# GRNFormer: Accurate Gene Regulatory Network Inference Using Graph Transformer

**DOI:** 10.1101/2025.01.26.634966

**Authors:** Akshata Hegde, Jianlin Cheng

## Abstract

**Motivation:** Deciphering gene regulatory networks (GRNs) from *single-cell* transcriptomics data remains a fundamental challenge in computational biology. It is hindered by data sparsity, high dimensionality, and the lack of scalable, generalizable inference models. To address this, we present GRNFormer, a generalizable graph transformer framework for accurate GRN inference from transcriptomics data across species, cell types, and platforms without requiring cell-type annotations or prior regulatory information.

**Results:** GRNFormer integrates a transformer-based gene expression encoder (Gene-Transcoder) with a variational graph autoencoder (GraViTAE) employing pairwise attention to jointly learn the representations of genes (nodes) and their co-expression relationships (edges). Leveraging TF-Walker, a transcription factor-anchored subgraph sampling strategy, it effectively captures gene regulatory interactions from either single-cell or bulk RNA-seq datasets. Benchmarking on standard datasets demonstrates that GRNFormer outperforms existing traditional and deep learning state-of-the-art methods in blind evaluations, achieving average Sampled Area Under the Receiver Operating Characteristic Curve (Sampled_AUROC) and Sampled Area Under the Precision-Recall Curve (Sampled_AUPRC) values between 0.90 and 0.98 as well as 0.87-0.98 average Sampled F1 score. The model robustly recovers both known and novel regulatory networks, including pluripotency circuits in human embryonic stem cells (hESCs) and immune cell modules in Peripheral Blood Mononuclear Cells (PBMCs). The architecture enables scalable, biologically interpretable GRN inference across various datasets, cell types, and species, establishing GRNFormer as a robust and transferable tool for network biology.

**Availability:** GRNFormer is available on GitHub (https://github.com/BioinfoMachineLearning/GRNformer); the version used in this work is archived on Zenodo (https://doi.org/10.5281/zenodo.18868395), with evaluation resources for reproducibility.

## 1 Introduction

Precise gene regulation is essential for normal cellular function, while its dysregulation is linked to diseases such as cancer, neurodegeneration, and developmental disorders. Understanding gene regulation at the network level-through gene regulatory networks ( GRNs)-can illuminate the interactions among genes and proteins that orchestrate cellular function. GRN inference enables the identification of key regulatory drivers and pathways, offering a foundation for mechanistic insights and therapeutic targeting (De Smet and Marchal 2010). However, inferring GRNs from high-throughput expression data remains challenging due to noise, high dimensionality, and limited sample sizes (Chen, Ning and Shi 2019).

Traditional approaches such as information-theoretic methods (e.g., ARACNE, CLR) (Margolin *et al*. 2006; Faith *et al*. 2007), or Bayesian networks (Shermin and Orgun 2009) can model gene dependencies but often require large datasets and face scalability limitations. Dynamic models, including Boolean networks and differential equation-based approaches (Hickman and Hodgman 2009; Lu *et al*. 2011), offer temporal insights but depend on time-resolved data, which is often lacking. The BEELINE benchmarking study (Pratapa *et al*. 2020) underscores these issues, showing that conventional statistical and shallow learning approaches struggle to generalize across diverse regulatory contexts. Recent approaches, particularly those based on deep learning, are capable of learning complex, non-linear patterns from high-dimensional data, making them better suited for inferring gene regulatory networks from RNA-Seq data (Huynh-Thu and Sanguinetti 2018; Yang, Fang and Shen 2019). In particular, transformers have revolutionized the perspectives of analyzing the input information through the “attention” mechanism (Vaswani *et al*. 2017), integrating global and local dependencies enabling improved GRN inference(Ma *et al*. 2021). There have been a few transformer-based GRN inference methods developed, such as STGRNs (Xu *et al*. 2023) and scGREAT (Wang *et al*. 2024). Complementing these, Graph Neural Networks (GNNs) provide a natural framework for modeling GRNs by learning over graph-structured data, effectively capturing both local and global regulatory patterns(Kipf and Welling 2016; Zhou *et al*. 2020). GeneLink is one such early link-prediction methods combining GNNs and attention.(Chen and Liu 2022)

However, most GRN inference methods are tailored to specific datasets, limiting their generalizability across species, cell types or conditions. Hence, there is a critical need for robust, transferable models that perform reliably beyond narrow contexts. In this study, we introduce GRNFormer, a graph deep learning framework developed to infer GRNs from either single-cell or bulk transcriptomic data with high accuracy and generalizability (Figure 1). Designed to operate across diverse cell types, species, and regulatory contexts, GRNFormer addresses key challenges in GRN inference such as context-specificity, data sparsity, and model transferability.

**Figure 1:**
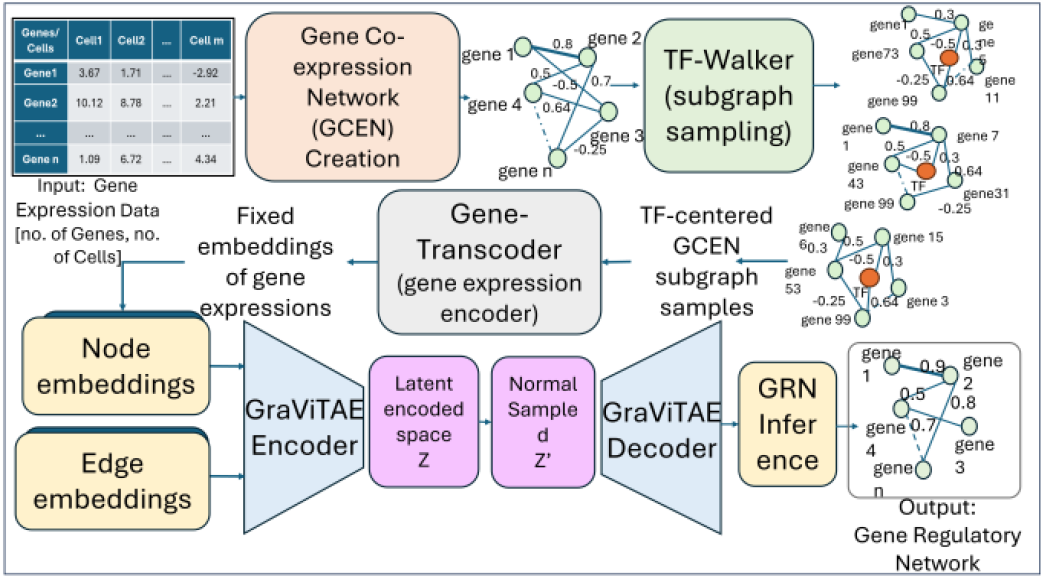
Overview of GRNFormer pipeline.

## 2 Methods

GRNFormer architecture comprises of three main components (Figure 1). First, TFWalker introduces a transcription factor-centered de-novo subgraph sampling approach that constructs localized gene co-expression subgraphs from a full gene co-expression network (GCEN) input, capturing the neighborhood context around each transcription factor(TF). This biologically informed strategy enhances the model’s ability to learn meaningful regulatory patterns by focusing on TF-driven structures in the gene expression space.

Second, we implement end-to-end representation learning through two key modules: the Gene Transcoder, a transformer-based embedding module that captures context-aware gene representations across diverse datasets; and GraViTAE (Graph Variational Transformer Autoencoder), which jointly encodes node and edge features to reconstruct gene regulatory feature representations.

Third, a dedicated GRN inference module integrates node- and edge-level representations to predict TF-target interactions. This inference strategy allows GRNFormer to generalize effectively across input sizes, data modalities, and regulatory frameworks, including both ChIP-seq-based and STRING(Szklarczyk *et al*. 2023)-derived networks.

### 2.1 Construction of gene co-expression network

To construct the gene co-expression network (GCEN), raw single-cell RNA-seq (scRNA-seq) expression data were first normalized using the inverse hyperbolic sine (Arc Sinh) transformation to stabilize variance across highly skewed expression profiles (Johnson and Krishnan 2022). Pairwise Pearson correlations were then computed between all genes across individual cells to quantify co-expression, and gene pairs with absolute correlation >0.1 were retained to form the GCEN. In this graph, nodes represent genes and edges denote co-expression relationships, forming a dense, high-dimensional network. The GCEN construction workflow is illustrated in Supplementary Method S1; Supplementary Fig. S1A.

### 2.2 TF-Walker - a de-novo subgraph sampling method

scRNA-seq data presents a fundamental challenge for machine learning: expression matrices are inherently high-dimensional and sparse, with expression values captured across thousands of genes and cells. However, these datasets often contain only a limited number of training examples. This imbalance results in under-constrained models, where the number of features far exceeds the number of samples needed for learning regulatory patterns.

To address these challenges, GRNFormer incorporates a biologically motivated subgraph sampling strategy, termed **TF-Walker (**Supplementary Figure S1B) that functions as a principled form of data augmentation. Instead of operating on the full GCEN, TF-Walker extracts local, transcription factor (TF)-centered subgraphs that capture meaningful co-expression contexts. To extract TF-centered subgraphs from the gene co-expression network, we iteratively process each transcription factor as a center node. For each TF, we first identify its direct neighbors (hop=1) in the network. If the total number of neighbors (T) is less than 99, we incrementally expand the neighborhood by increasing the hop distance (hop+1) and recursively collecting neighbors at each subsequent hop until T reaches or exceeds 99. During training, we fixed the subgraph size to 100 nodes (one TF node and 99 neighbors), which balances capturing meaningful regulatory context, minimizing overlap among subgraphs, and computational efficiency. If the neighborhood size exceeds T, we do single random selection of 99 neighbors. Once the target neighborhood size is achieved, we select all edges present between the nodes within the selected neighborhood, ensuring comprehensive connectivity representation. Subsequently, we extract all node and edge features associated with the selected subgraph, including binary TF identities, co-expression weights (positive or negative correlations). This process generates a TF-centered subgraph for each transcription factor, where the TF serves as the central hub connected to up to 99 neighboring genes through their co-expression relationships, thereby capturing the local regulatory context surrounding each TF within the broader network topology. This procedure is repeated for all identified transcription factors in the network, resulting in a collection of TF-specific subgraphs that preserve the structural and functional characteristics of each TF’s regulatory neighborhood while maintaining computational tractability through the size constraint. The TF-Walker sampling algorithm is shown in Supplementary Fig. S1B and formalized in Supplementary Method S2. In contrast, during inference, TF-Walker deterministically extends to all available neighbors of a given TF, traversing the GCEN in a sequential and exhaustive manner to generate subgraphs. This ensures that regulatory predictions are based on the full local topology of the expression network, without arbitrary truncation. Additional details on the inference strategy are provided in the GRN Inference section.

### 2.3 Ground Truth Network Sampling and Feature Representation

To supervise GRNFormer training, we constructed comprehensive ground truth regulatory networks by integrating three validated sources: cell-type-specific ChIP-seq data, non-specific ChIP-seq networks, and STRING-derived protein-protein interactions. This multi-source integration enhances biological coverage and supports cross-species generalization. For each TF-centered subgraph generated by TF-Walker, a corresponding labeled regulatory subgraph was created by identifying all nodes (genes) in the subgraph and extracting known regulatory interactions among them from the global ground truth network. If no regulatory edges existed between nodes, the subgraph was discarded to ensure only biologically informative samples were used for training. This approach guarantees that the model learns from functionally relevant regulatory relationships, improving inference reliability.

Each subgraph is represented as a graph of 100 genes (nodes), where the features of each node consist of the expression values of the gene across all cells along with a binary indicator denoting TF identity (i.e., if a gene is a TF). Edge features are defined by pairwise Pearson correlation values computed from the full dataset, capturing global co-expression structure. Gene expression of each subgraph undergoes cellwise z-score normalization across genes. This normalization mitigates information leakage by preventing memorization of gene identities/patterns across subgraphs while preserving local expression dynamics. While the edge weights reflect broader transcriptional associations, the node-level gene expression inputs provide localized information contextualized at the subgraph level.

### 2.4 Gene-Transcoder: Transformer encoder for gene expression representation learning

Figure 2A illustrates the architecture of the Gene-Transcoder, a transformer-based encoder designed for gene expression representation learning, which is the first stage in training GRNFormer. It is designed to address variability in scRNA-seq datasets, where the number of cells and expression ranges can differ substantially across experiments, making it challenging for conventional models to learn generalizable patterns. To tackle this issue, the Gene-Transcoder processes gene expression data to produce fixed, representative embeddings that capture essential biological information.

**Figure 2:**
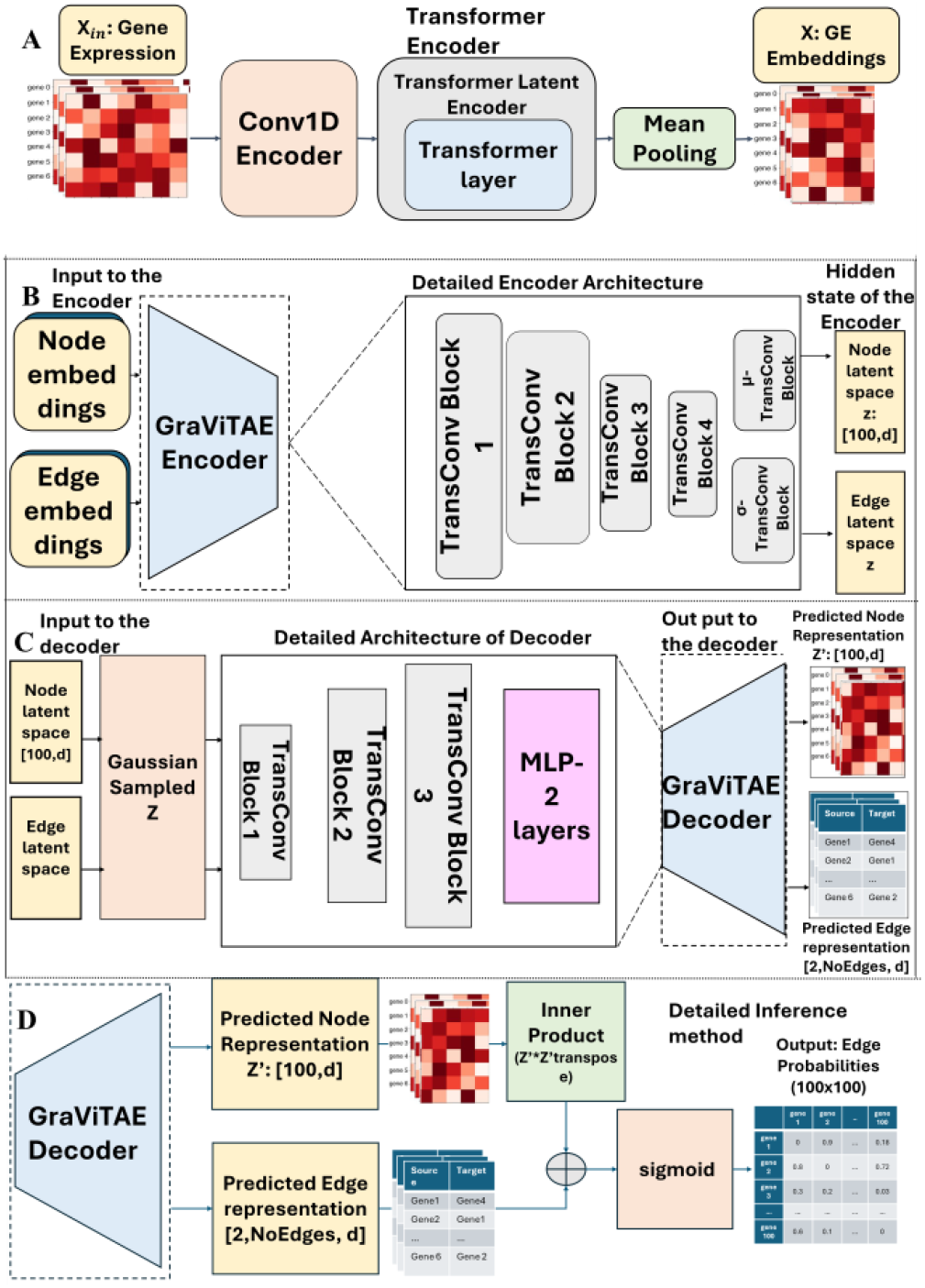
The four key sequentially connected components of GRN-Former. **(A)** Architecture of Gene Transcoder for fixed embeddings of gene expression. **(B)** Detailed architecture of GraViTAE Encoder. **(C)** Detailed architecture of GraViTAE variational decoder. **(D)** Edge probability prediction of GRN inference module

Gene-Transcoder begins with an input gene expression matrix (*batch* × *100 genes* × *no_of_cells*+*1*), where the additional channel corresponds to the TF-identity flag appended along the cell/feature dimension during feature extraction. The 100-gene input dimension used in Gene-Transcoder corresponds to the TF-Walker subgraph size during training. A 1D convolutional layer processes the gene expression matrix. This convolution utilizes a sliding kernel of size 1, and stride 1 along the cell axis to capture local patterns of gene expression and TF-related signal across cells. These features are then passed to a transformer encoder of single layer, where a 4-head attention mechanism is employed. This mechanism helps the model to jointly capture local expression dependencies among nearby genes and global relationships that span distant genes or cell populations and produces embeddings that represent gene expression data in an embedding feature space.

The transformer outputs are averaged across cells via mean pooling to produce fixed 64-dimensional embeddings (*batch* × *100* × *64*), providing a compact yet information-rich representation for each gene. These embeddings are invariant to dataset-specific variation, enabling effective generalization across species, cell types, and experimental conditions. Detailed mathematical algorithm for the Gene-Transcoder is provided in in Supplementary Method S3; Supplementary Fig. S2. By transforming variable-length, noisy expression profiles into fixed, context-aware embeddings, this module establishes a robust foundation for downstream GRN inference.

### 2.5 Graph Variational Transformer Autoencoder (GraViTAE)

Following Gene-Transcoder is the Graph Variational Transformer Auto-encoder (GraViTAE), that forms the core of GRNFormer. GraViTAE is a supervised variational graph transformer autoencoder composed of an encoder–decoder pair operating on TF-centered gene co-expression subgraphs. It integrates fixed-length node embeddings from the Gene-Transcoder with co-expression edge weights for joint message passing.

As shown in Figure 2B, the GraViTAE encoder, applies stacked Transformer Convolution (TransConv) blocks to capture both local (e.g., between closely related genes) and global gene-gene interactions (e.g., co-expression relationships across a subgraph). Each block combines multi-head attention (4-head) with feedforward layers to update the node and edge features jointly. The GraViTAE encoder produces the parameters of a Gaussian latent distribution for each TF-centered subgraph. Let *z*^(*v*)^denote the latent embedding of the TF-centered subgraph for transcription factor *v*. Given input features x, the encoder outputs a mean and standard deviation vector, μ_ϕ_(x)and σ_ϕ_(x), defining the approximate posterior

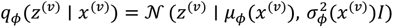

where *ϕ*: parameters of the encoder network. 𝒩(⋅∣ *μ*, σ^2^): multivariate normal distribution with mean *μ* and covariance 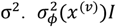: diagonal covariance matrix, where *I* is the identity matrix. To regularize the latent space and promote generalization, the encoder estimates distributions over latent variables by computing both the mean (μ) and variance (σ^2^) of node and edge embeddings. These parameters are used to sample continuous latent embeddings via a Gaussian reparameterization trick.

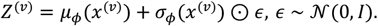

Where, *Z*^(*v*)^: latent embedding sampled for the TF-centered subgraph. This variational formulation enables uncertainty modeling in noisy single-cell expression data, while the joint encoding of expression and co-expression features ensures biologically grounded, context-aware representations. The GraViTAE decoder (Figure 2C) reconstructs node and edge embeddings from the sampled latent space of the encoder. Using these samples, the decoder applies additional TransConv blocks followed by a light-weight multilayer perceptron (MLP) to produce two outputs: updated node embeddings *Z*′ representing each gene’s regulatory identity and edge attentions, indicating the strength of potential regulatory interactions. These outputs serve as intermediate proxy representations to guide the down-stream GRN inference.

The TransConv block (Supplementary Figure S3A) underpins both GraViTAE encoder and decoder modules (Figure 2B and 2C). At the heart of the TransConv block lies the *Transformer Convolution* layer. It extends transformer-based graph convolution (Shi *et al*. 2020) by incorporating edge attributes such as co-expression into its pairwise attention computations, along with node features, enabling context-aware message passing. The pairwise attention coefficient *a*_*ij*_ between genes i and j is defined as

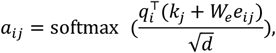

where *q*_*i*_ = *W*_*q*_*x*_*i*_, *k*_*j*_ = *W*_*k*_*x*_*j*_, and *e*_*ij*_is the edge feature. The pairwise attention details, message-passing, and update steps implemented through the Transformer Convolution layer are provided in Supplementary Method S4 and Supplementary Fig. S3B. Each block includes multi-head pairwise attentions with residual connections, feed-forward layers, and nonlinear transformations, followed by batch normalization. We employ Leaky ReLU activations to preserve gradient flow and capture negative co-expression patterns often associated with repressive regulation that standard ReLU activations tend to suppress (Xu et al. 2020)(Xu et al. 2020). Implemented in *PyTorch Geometric*, this architecture effectively models the combinatorial and context-dependent nature of transcriptional regulation.

### 2.6 Adjacency Matrix Reconstruction and GRN Inference

The final step of GRNFormer reconstructs a probabilistic adjacency matrix representing the likelihood of regulatory interactions between genes. This is achieved by combining inner products of node embeddings with pooled edge features, followed by a sigmoid activation to produce gene– gene regulatory interaction probabilities. (Figure 2D). Detailed steps of adjacency matrix reconstruction is described in Supplementary Method S5. This subgraph level inference captures modular gene regulation, while a dedicated aggregation strategy integrates predictions across subgraphs to reconstruct a coherent global GRN (see GRN Inference Section for details).

### 2.7 Datasets and Preprocessing

#### BEELINE data

To train and evaluate GRNFormer in biologically diverse settings, we used single-cell RNA-seq datasets curated by BEELINE, a comprehensive benchmarking framework for GRN inference. These datasets span two species (human and mouse), seven cell types, and include three types of regulatory ground truth networks, offering a broad foundation for testing model generalization across cell types, regulatory contexts, species, and evolutionary backgrounds.

Our analysis focused on all seven cell-type representative datasets: mouse hematopoietic stem cells (mHSC-E, mHSC-GM, mHSC-L), embryonic stem cells (mESC), dendritic cells (mDC), and human hepatocytes (hHep) and embryonic stem cells (hESC). Together, they reflect a wide range of biological processes from early development and lineage commitment to immune regulation and tissue-specific differentiation. All expression data were preprocessed using BEELINE’s standardized workflow, including log transformation, gene filtering, and normalization, ensuring consistency and cross-dataset comparability. Each expression dataset was paired with three regulatory reference networks (i) cell-type-specific TF– target interactions from experimental ChIP-seq, (ii) non-cell-type-specific ChIP-seq networks, and (iii) functional protein interaction networks from STRING. The combined use of regulatory evidence enabled robust bench-marking of GRNFormer’s predictions in both precise and broad regulatory contexts.

#### DREAM5 Challenge and PBMC Datasets

We also applied GRNFormer, pretrained on the BEELINE single cell data to bulk RNA-seq datasets from the DREAM5 challenge (Marbach et al. 2012)(Marbach et al. 2012), that includes *Escherichia coli* and *Saccharomyces cerevisiae* data. These well-characterized prokaryotic and eukaryotic networks, with gold-standard ChIP-derived regulatory maps, evaluated model’s cross-species transferability and robustness to different transcriptomic data types. To further test generalizability, we conducted zero-shot inference using the 10x Genomics PBMC 3k dataset (Genomics 2018)(Genomics 2018), which lacks any cell-type labels or prior regulatory annotations. This allowed us to assess GRNFormer’s ability to recover meaningful regulatory structure de novo.

### 2.8 Training

To assess the generalizability of GRNFormer, we adopted a cross-lineage training and evaluation strategy. The model was trained on subgraphs derived from five datasets - human embryonic stem cells (hESC), human hepatocytes (hHep), mouse dendritic cells (mDC), and two mouse hematopoietic stem cell subtypes (mHSC-GM, mHSC-E) - while holding out mouse embryonic stem cells (mESC) and an additional mHSC subtype (mHSC-L) for blind testing. For each dataset, we first constructed full gene co-expression networks (GCENs) and then applied TF-Walker subgraph sampling to generate localized training instances. Ground-truth regulatory interactions were aggregated from three sources - cell-type-specific ChIP-seq, non-specific ChIP-seq, and STRING networks - into a unified training label set, allowing the model to learn from heterogeneous regulatory modalities.

Training on shuffled subgraphs from diverse datasets encouraged the model to capture transferable transcriptional patterns rather than overfitting to dataset-specific profiles. The Gene-Transcoder module encodes variable-length gene expression profiles of subgraph genes into fixed-length node embeddings of dimension 64. These node embeddings, along with co-expression values used as edge features, are passed to the encoder implemented as a part of variational graph transformer autoencoder (GraViTAE). The encoder comprises four Transformer Convolution (TransConv) blocks, each with 4 attention heads, and projects the input features into a latent space of dimension 16.

The sampled latent node and edge representations are then passed through the GraViTAE decoder, which consists of three additional TransConv blocks followed by two fully connected (MLP) layers, reconstructing interaction-specific representations. These intermediate outputs are aggregated and transformed into probabilistic edge predictions used to infer the GRN. GRNFormer was trained for 100 epochs with a batch size of 8, using a composite loss function that combines binary cross-entropy (BCE) for predicting regulatory edge presence with Kullback–Leibler (KL) divergence to regularize the variational latent space. The binary cross-entropy reconstruction loss is

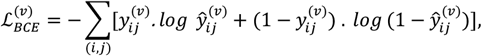

which measures the discrepancy between the predicted probabilities 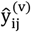 and the true binary labels 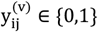. To regularize the latent representation z^(v)^, we include the Kullback–Leibler (KL) divergence between the approximate posterior q_ϕ_(z^(v)^ ∣ x^(v)^)and a standard normal prior p(z). The normalized KL term is

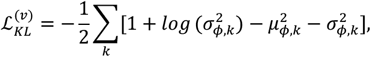

where μ_ϕ,k_and 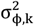 are the mean and variance of the k-th component of the latent vector z^(v)^.

The total loss combines the reconstruction and regularization terms as

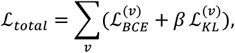

where β is a scaling factor (e.g., 0.01or 1/#genes in subgraph) controlling the strength of KL regularization.

The training objective is framed as a binary classification task: identifying whether an edge between any two genes within a subgraph corresponds to a known regulatory interaction. Detailed loss functions and variational formulation are also explained in Supplementary method S6 along with Evidence Lower Bound (ELBO) formalization of the model. The model used Adam optimizer (initial learning rate 0.001) with a ReduceLROnPlateau scheduler and early stopping. L1 regularization was applied to prevent overfitting.

To address extreme class imbalance-typical of GRN inference where true regulatory edges are sparse, we employed dynamic negative sampling during training. For each subgraph, we matched the number of negative (non-regulatory) edges to the number of positives, ensuring balanced and effective learning across highly imbalanced edge distributions. Importantly, the model was trained without any explicit knowledge of species, cell type, or regulatory context, and therefore it can be applied across diverse species, cell types, and regulatory contexts. All subgraphs were randomly shuffled prior to training, forcing the model to rely solely on expression-derived contextual signals. GRNFormer is trained on 2 GPUs of NVDIA A10, on national GPU clusters of Nautilus server. Table 1 consolidates the essential configuration settings for Gene-Transcoder, GraViTAE, and TF-Walker, as well as the training procedure.

**Table 1.** Summary of key architectural and training hyperparameters used in GRNFormer.

| Module | Key settings |
| --- | --- |
| Gene-Transcoder | 1D conv (kernel = 1, stride = 1); single-layer transformer encoder (4 heads); output embedding = 64 dims; input = 100 genes × (#cells + TF flag). |
| GraViTAE (Encoder/Decoder) | 3 TransConv blocks (encoder) + 3 TransConv blocks (decoder); 4 attention heads per block; hidden size = 64; latent mean/variance dim = 16. |
| TF-Walker | Training subgraph size = 100 genes; co-expression threshold = 0.1; |
| Training | Adam (lr = 0.001); ReduceLROnPlateau; batch size = 8; epochs = 100; L1 regularization; loss = BCE + normalized KL divergence ($\beta = 1 / \text{\#genes\_in\_subgraph}$). |

### 2.9 GRN Inference and Evaluation on Test Datasets

During inference, GRNFormer employs an extended version of the TF-Walker strategy to generate high-coverage subgraphs for each test dataset. Unlike training, where 99 neighbors are sampled randomly, inference subgraphs are constructed sequentially to include all co-expressed neighbors of each TF, ensuring maximal gene and interaction coverage. Multiple subgraphs may contain overlapping genes; however, each subgraph is independently z-score normalized, treating recurring genes as context-specific instances to enhance robustness. The deterministic expansion of TF-Walker algorithm used during inference stage is illustrated in Supplementary Method S7; Supplementary Figure S4.

Predicted interactions from all subgraphs are aggregated to reconstruct the full GRN, with a focus on TF–gene and TF–TF interactions that form the regulatory core. Within each subgraph, interactions with predicted probability >0.3 were retained to capture both confident and potentially weaker but biologically relevant regulatory relationships, recognizing that gene regulation often involves subtle, low-strength interactions. TF-initiated edges were prioritized during aggregation. Each interaction was assigned a weighted score based on prediction probability, TF involvement, and frequency across subgraphs. These scores were summed, averaged, and min–max normalized to produce the final adjacency matrix, with entries representing the likelihood of regulatory relationships between gene pairs. We evaluated GRNFormer using standard classification metrics-area under the receiver operating characteristic curve (AUROC), the area under the precision-recall curve (AUPRC), F1 score, precision, recall, accuracy and early precision (EPR),-each offering a complementary insight into performance under class imbalance. Metric definitions are provided in Supplementary Methods S8. Context-specific evaluations were conducted on the BEELINE cross-cell-type test dataset (mESC, mHSC-L), focusing on biologically relevant gene subsets (e.g., top 500–1,000 most variable genes, with or without TFs) to assess model robustness under constrained feature spaces.

Given the absence of annotated true negatives in GRN benchmarks, we adopted an enhanced negative sampling strategy based on bootstrapping. Following practices used in GRLGRN (Wang et al., 2025), CNNC, and scGREAT, we randomly sampled gene pairs not found in the ground-truth network as negatives. For each test dataset, we paired known TF–target interactions (positives) with an equal number of randomly sampled negatives. This process was repeated over 100 bootstrap iterations, reducing sampling bias and providing statistically robust performance estimates. In each iteration, classification metrics were computed based on predicted probabilities and the combined set of positives and sampled negatives. Final scores were averaged across all iterations. We primarily report AUROC and AUPRC, in benchmarks which are most reliable in evaluating performance on sparse, imbalanced regulatory networks. When we calculate AUROC and AUPRC using the bootstrapped negative samplings, we report them as Sampled_AUROC and Sampled_AUPRC and full matrix evaluations are reported as Full test-set AUROC/AUPRC.

Hyperparameter sensitivity analyses for the GCEN co-expression threshold and TF-Walker subgraph size were performed at the inference stage (Supplementary Method S11). Supplementary Fig. S5 shows that GCEN thresholds of 0.1–0.3 yield stable performance, and Supplementary Fig. S6 and Sup. Table S4 indicate that TF-Walker subgraph sizes of 100–200 genes are consistently optimal. In our main experiments, we used GCEN = 0.1 and a subgraph size of 100, and these parameters are configurable in the GRNFormer codebase for application-specific tuning. Guidelines for selections are provided in Supplementary Method S11 C.

## 3 Results

### 3.1 High-accuracy Inference of GRNs Across Regulatory Contexts

To evaluate the generalizability and contextual sensitivity of GRNFormer, we conducted a comprehensive benchmarking study using the BEELINE suite of single-cell gene expression datasets. Our goal was to assess whether GRNFormer could robustly infer GRNs across species boundaries, cell identities, and diverse regulatory contexts without relying on prior knowledge.

GRNFormer was trained to infer GRNs solely from gene and transcription factor (TF) expression profiles, without providing information about gene names, cell type, species, or regulatory context during training or evaluation. This design encourages inductive generalization across diverse biological settings. During evaluation, global GRNs were reconstructed by aggregating regulatory interactions inferred from locally sampled, TF-centered co-expression subgraphs.

Training and internal evaluation were performed on five cell types (hESC, hHep, mDC, mHSC-E, and mHSC-GM). Although individual genes may appear in both phases, we performed localized z-score normalization across genes for each cell in every subgraph, ensuring that the model treats each instance independently and preventing memorization. To rigorously assess generalization and guard against information leakage, we conducted a cross-dataset evaluation on two held-out cell types - mESC and mHSC-L - not seen during training.

Table 2 summarizes GRNFormer’s average performance on these two withheld cell types (mESC and mHSC-L) unseen in the training across regulatory settings, reporting Sampled_AUROC(SAUROC), Sampled_AUPRC(SAUPRC), Sampled - F1 score, precision, recall, and accuracy for each species-cell type-regulatory context combination. The results of the internal evaluation on the other five cell types are presented in Supplementary Table S1. During the cross-dataset evaluation, we evaluated GRNFormer’s performance on subsets of the dataset of each cell type (mESC and mHSC-L) containing the top 500 and 1,000 highly variable genes, both with and without the most significantly varying TFs. The test subsets were prepared according to BEELINE’s standard data preprocessing and evaluation protocols.

**Table 2.** Summary of evaluation metrics of GRNFormer across BEELINE test dataset.

| Network Type | Cell Type | Gene sets | #Genes | #True Positives | SAUROC | SAUPRC | Precision | Recall | F1Score | Accuracy |
| --- | --- | --- | --- | --- | --- | --- | --- | --- | --- | --- |
| Non-Cell-Type specific ChIP-Seq | mESC | 1000 | 1000 | 852 | 0.965 | 0.938 | 0.925 | 0.998 | 0.96 | 0.959 |
|  |  | 500 | 500 | 264 | 0.97 | 0.953 | 0.92 | 0.999 | 0.958 | 0.956 |
|  |  | TF+1000 | 1000 | 853 | 0.965 | 0.939 | 0.926 | 0.998 | 0.961 | 0.959 |
|  |  | TF+500 | 500 | 264 | 0.965 | 0.943 | 0.918 | 0.999 | 0.957 | 0.955 |
|  | mHSC-L | 1000 | 1000 | 589 | 0.984 | 0.974 | 0.956 | 0.999 | 0.978 | 0.977 |
|  |  | 500 | 500 | 109 | 0.981 | 0.966 | 0.956 | 1 | 0.977 | 0.977 |
|  |  | TF+1000 | 1124 | 771 | 0.979 | 0.964 | 0.945 | 0.999 | 0.972 | 0.971 |
|  |  | TF+500 | 624 | 467 | 0.942 | 0.903 | 0.886 | 0.859 | 0.872 | 0.874 |
| Cell Type Specific ChIP-Seq | mESC | 1000 | 1000 | 5499 | 0.967 | 0.942 | 0.93 | 0.998 | 0.963 | 0.962 |
|  |  | 500 | 500 | 1913 | 0.968 | 0.944 | 0.923 | 0.999 | 0.959 | 0.958 |
|  |  | TF+1000 | 1000 | 5500 | 0.968 | 0.943 | 0.93 | 0.999 | 0.963 | 0.962 |
|  |  | TF+500 | 500 | 1913 | 0.967 | 0.943 | 0.923 | 0.999 | 0.96 | 0.958 |
|  | mHSC-L | 1000 | 1000 | 1325 | 0.979 | 0.959 | 0.956 | 0.999 | 0.978 | 0.977 |
|  |  | 500 | 500 | 1295 | 0.979 | 0.954 | 0.96 | 0.999 | 0.98 | 0.979 |
|  |  | TF+1000 | 1124 | 7746 | 0.976 | 0.954 | 0.949 | 1 | 0.974 | 0.973 |
|  |  | TF+500 | 624 | 5182 | 0.943 | 0.898 | 0.89 | 1 | 0.942 | 0.938 |
| STRING | mESC | 500 | 500 | 264 | 0.957 | 0.928 | 0.916 | 0.996 | 0.954 | 0.952 |
|  |  | 1000 | 1000 | 661 | 0.963 | 0.937 | 0.926 | 1 | 0.961 | 0.960 |
|  |  | TF+1000 | 1000 | 661 | 0.964 | 0.937 | 0.925 | 0.991 | 0.957 | 0.955 |
|  |  | TF+500 | 500 | 263 | 0.959 | 0.931 | 0.917 | 0.992 | 0.953 | 0.951 |
|  | mHSC-L | 500 | 500 | 14 | 0.982 | 0.973 | 0.957 | 1 | 0.977 | 0.976 |
|  |  | 1000 | 1000 | 537 | 0.991 | 0.982 | 0.956 | 1 | 0.977 | 0.977 |
|  |  | TF+1000 | 1124 | 114 | 0.981 | 0.972 | 0.941 | 1 | 0.969 | 0.968 |
|  |  | TF+500 | 624 | 67 | 0.944 | 0.913 | 0.89 | 0.855 | 0.872 | 0.874 |

As shown in Table 2, GRNFormer consistently achieves high accuracy across all the subsets spanning different network types (non-specific-ChIP-Seq network, cell-type-specific ChIP-Seq network, and STRING-based network), cell types, and gene sets. High Sampled_AUROC and Sampled_AUPRC scores between ~0.90 and 0.98 as well as high Sampled F1 scores between 0.87 and 0.98, indicates robust, accurate, generalized GRN inference across diverse conditions.

### 3.2 GRNFormer Outperforms State-of-the-art GRN Inference Methods across Single-cell Benchmarks under Blind Evaluation

To systematically evaluate the performance of GRNFormer, we conducted multiple comprehensive benchmarking analyses against nine widely used gene regulatory network (GRN) inference methods: with five deep-learning methods, CNNC(Yuan and Bar-Joseph 2019)(Yuan and Bar-Joseph 2019), GNE(Kc et al. 2019)(Kc et al. 2019), GNNLink(Mao et al. 2023)(Mao et al. 2023), STGRNS, and scGREAT and four traditional methods, LEAP(Jiang and Neapolitan 2015)(Jiang and Neapolitan 2015), PIDC(Chan, Stumpf and Babtie 2017)(Chan, Stumpf and Babtie 2017), PPCOR(Kim 2015)(Kim 2015), SINCERITIES(Papili Gao et al. 2018)(Papili Gao et al. 2018). Supplementary Table S2 provides the brief summary of each method used, and Supplementary method S9 provides details on benchmark setup. Performance was assessed on three independent gold-standard reference networks: (i) non-cell-type specific ChIP-seq, (ii) cell-type-specific ChIP-seq, and (iii) functional interaction networks from STRING. For each method and evaluation setting, we computed, Full test-set AUROC/AUPRC, Full test-set Early Precision (EPR) and Sampled_AUROC/AUPRC. Gene selection followed the BEELINE protocol, using four standard configurations: (1) top 500 highly variable genes (500), (2) top 1,000 variable genes (1000), (3) TFs + top 500 (TF500), and (4) TFs + top 1,000 (TF1000) variable genes. While BEELINE primarily targets unsupervised GRN inference, our benchmarking includes both supervised and unsupervised methods; therefore, we extend the evaluation framework through a unified clean negative pool to enable fair comparison across learning paradigms.

GRNFormer was trained once on general dataset spanning five cell types (hESC, hHep, mDC, mHSC-E, mHSC-GM) and evaluated on two unseen cell types (mESC and mHSC-L) without task-specific fine-tuning, simulating a stringent, cross datasets generalization scenario. In contrast competing deep learning methods were trained and evaluated on within-dataset splits of each test cell type, following protocols described in their original publications.

This less stringent evaluation strategy was applied to the nine state-of-the-art (SOTA)methods, because, as described in their original publications, most of these models were optimized for within-dataset performance and did not perform cross-cell-type generalization. For consistency and fairness, we reproduced each competing method using the same training–validation splits but retained their originally reported training procedures. In all evaluations, only positive regulatory interactions were split into training, validation, and test sets. For sampled metrics, negative edges were bootstrapped in a 1:1 ratio from a clean negative evaluation pool, whereas full test-set AUROC and AUPRC were computed using the entire clean negative pool. This clean pool is constructed by removing all training/validation/test positives, all training/validation negatives, and self-loops from the full gene–gene space, ensuring that none of the negative edges used for evaluation were seen during any of the model’s training. The construction of the clean negative evaluation pool is described in Supplementary Method S10.

To evaluate performance under realistic extreme class imbalance, we first performed a full-matrix evaluation using the positive test set and complete clean negative evaluation pool which served as unseen negatives (see Supplementary Method S10 for detailed protocol, Supplementary Figure S8B for test-positives-to-negative edge counts).

Figure 3 presents the comparison heatmaps of Full test-set AUROC and Full test-set AUPRC across all methods and test datasets under this standardized full-matrix evaluation. Even though GRNFormer was tested on cell types it had never encountered during training while several competing methods were trained and tested within the same cell type, GRNFormer consistently achieves the highest or near-highest performance across most of the datasets. In Full test-set AUROC (Figure 3A), GRNFormer maintains strong performance in both non–cell-type-specific and STRING network settings (Full test-set AUROC >0.8 21/24 datasets, with particularly robust values in the mESC STRING benchmark and the Non - ChIP-seq TF500/TF1000 gene sets(Full test-set AUROC >0.9). In terms of Full test-set AUPRC (Figure 3B), GRNFormer shows clear advantages in precision–recall behavior, especially on non-cell-type-specific and cell-type-specific ChIP-seq datasets, where many competing methods experience substantial degradation. Traditional statistical approaches such as PPCOR, and SINCERITIES continue to perform poorly in these settings, reflecting their limited ability to generalize. Among the external supervised competing methods GNE emerges as the strongest, achieving the most competitive Full test-set AUROC and Full test-set AUPRC whereas LEAP emerges as a strongest competing unsupervised method across datasets under the stringent full-matrix setting. GNNLink also performs well on several datasets on Full test-set AUROC, but its Full test-set AUPRC varies considerably across evaluation settings, reflecting its greater sensitivity to dataset characteristics. In contrast, GRNFormer surpasses all SOTA methods-including GNE-on the majority of datasets, with especially strong consistency in Full test-AUPRC, where precise identification of true regulatory relationships is most critical (See Sup. Results S1B for detailed analyses). To further evaluate early-ranking performance, we computed Early Precision (EP) across multiple cutoffs (Supplementary Results S1.C, Sup. Figs. S10 and S11). GRNFormer demonstrates consistently strong and stable recovery of true regulatory interactions at cutoffs (EP@1000, EP@2000, and EPR@# ground-truth(gt)),with GNE being equally competitive at lower cutoffs EP@100,EP@gt, and GRNFormer outperforms other competing methods across datasets with the average win rate of 84% (GRNFormer ≥ other methods). Win rate (Supplementary Fig. S11A) and rank frequency (Supplementary Fig. S11B) further confirm GRNFormer’s robust prioritization of biologically relevant regulatory edges.

**Figure 3:**
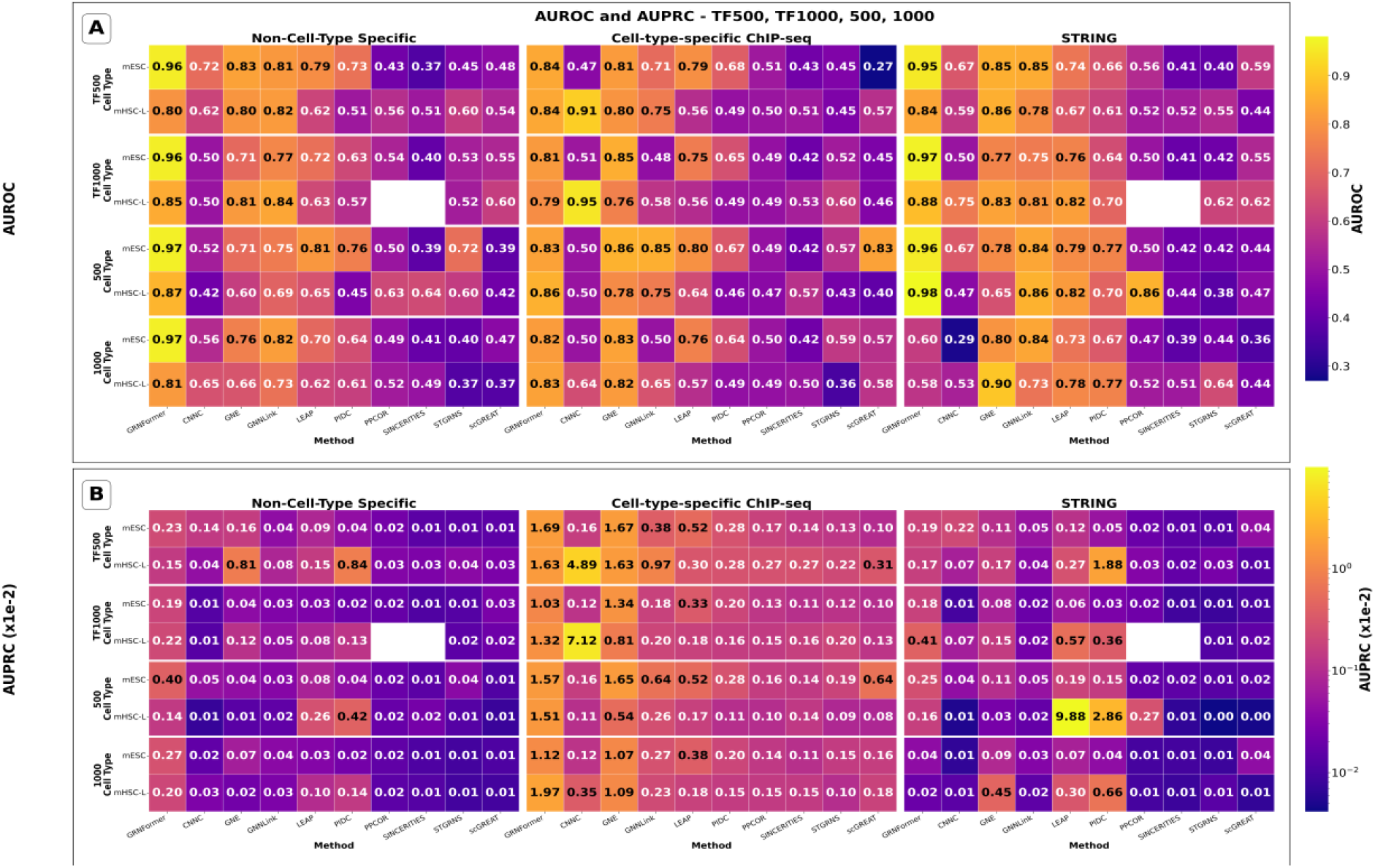
Comparison of GRNFormer and nine state-of-the-art methods in various test datasets. **(A)** Full test-set AUROC scores for 10 GRN inference methods evaluated on the BEELINE test set using three gold-standard benchmark GRNs: non-cell-type-specific ChIP-seq, cell-type-specific ChIP-seq, and STRING protein-protein interaction network. GRNFormer consistently ranks among the top performers across all evaluation scenarios. **(B)Full testset** AUPRC scores for the same evaluation settings. For readability, values are displayed after factoring out **×1e−2**.

To complement the full-matrix evaluation we additionally computed balanced sampled metrics following common supervised benchmarking practice. Specifically, we perform 100 iterations of 1:1 bootstrapped negative sampling from the same clean evaluation pool, pairing each test positive with a randomly drawn unseen negative edge. Because these metrics are derived from balanced sampling rather than the full edge space, they are explicitly reported as Sampled AUROC and Sampled AUPRC as Supplementary Fig. S7A–B.

(See Supplementary Method S10 for detailed protocol, detailed Results in Supplementary Results S1.A). These sampled evaluation and full matrix robustness evaluations(Supplementary Results S1A.B.C; Supplementary Fig. S7A–B,S8 & S9,S10&S11) confirm that GRNFormer is the top-performing method across more than half of datasets, with GNE emerging as the strongest competing method under the both sampled and stringent full-matrix setting. These results demonstrate that GRNFormer reliably infers high-confidence GRNs in a data-agnostic manner and achieves state-of-the-art accuracy under a stringent evaluation framework in which all methods are tested against the same unseen positives and negatives. This robustness across diverse benchmarks underscores GRNFormer’s practical utility for single-cell regulatory genomics.

### 3.3 Ablation Study: Dissecting GRNFormer’s Architecture

To assess the contribution of each architectural component in GRNFormer, we performed a systematic ablation study using a cross-lineage blind evaluation. The model was trained on five diverse expression datasets (hESC, hHep, mDC, mHSC-E, mHSC-GM) and blindly evaluated on two unseen cell types (mESC and mHSC-L) to rigorously test generalization across species and lineages. Ablation results shown in Supplementary Results S2 and Supplementary Table S3 demonstrate that the full GRNFormer model-comprising the TF-Walker subgraph sampler, Gene Transcoder, Transformer Convolution (TransConv) block, and GraViTAE decoder - achieved the highest performance (Sampled_AUROC = 0.97, Sampled_AUPRC = 0.95), confirming its robustness across biological contexts. The study reveals that TF-Walker effectively localizes transcriptional context, the Gene Transcoder and TransConv Block enrich representation learning, and the decoder improves accuracy in predicting regulatory interactions. Together, these components contribute to improved GRN reconstruction performance across diverse transcriptomic contexts.

### 3.4 Cross-Species Generalization on DREAM5 Bulk RNA Seq Datasets

To evaluate GRNFormer’s ability to generalize beyond mammalian single-cell data, we conducted a blind assessment using bulk RNA-seq datasets from *Escherichia coli* and *Saccharomyces cerevisiae* curated by the DREAM5 challenge. These benchmarks span distinct regulatory architectures-prokaryotic and unicellular eukaryotic-and are derived from high-confidence ChIP-based GRNs. GRNFormer, trained solely on human and mouse single-cell RNA-seq, was tested on these microbial bulk datasets without using species-specific annotations or prior regulatory knowledge. This evaluation addressed two core questions: (1) Can GRNFormer generalize across species with divergent transcriptional logic? (2) Does it retain accuracy when applied to bulk RNA-seq without retraining?

Applied directly to DREAM5 expression matrices, GRNFormer achieved Sampled_AUROC scores of 0.979 (*E. coli*) and 0.977 (*S. cerevisiae*), with Sampled_AUPRCs of 0.955 and 0.957, respectively, despite GRNFormer never seeing bulk or microbial data during training. Precision-Recall and Receiver Operating Characteristic curves are shown in Supplementary Results S3;Supplementary Figure S12. To ensure reliability, we used 100 rounds of bootstrapped negative sampling (1:1 ratio) and averaged metrics to mitigate variance. These findings confirm GRNFormer’s broad utility: it captures regulatory principles conserved across species, performs robustly across transcriptomic modalities, and scales up to genome-wide data without retraining. Its robust performance on DREAM5 underscores its potential as a general-purpose GRN inference engine, particularly in settings where ground-truth networks are limited or unavailable.

### 3.5 Scalability and Robustness of GRNFormer across Gene Set Size, Dataset Complexity, and Species

We evaluated the computational and biological scalability of GRNFormer using a diverse set of expression datasets varying in gene number, cell type, species, and regulatory context. Our analysis included all single-cell RNA-seq datasets from the BEELINE suite and bulk RNA-seq datasets from *E. coli* and yeast provided by the DREAM5 challenge. As shown in Figure 4C, inference runtime scaled predictably with the number of input genes (ranging from ~500 to ~5,900 genes), increasing smoothly on a logarithmic scale despite high dimensionality. Notably, this trend held across a wide range of network densities, reflecting the efficiency of the TF-Walker sampling strategy and the fixed-size subgraph processing pipeline. To assess computational scalability, all inference experiments were conducted on a single NVIDIA A100 GPU. For inference, the maximum peak memory usage was less than 2,500 MB, while the final memory usage after completion remained below 1,000 MB. For 4,911 genes, inference required 1,732.56 s, with a peak memory usage of 1,074.54 MB and a final memory usage of 380.87 MB. For 5,910 genes, inference took 4,512.08 s, with a peak memory usage of 2,406.18 MB and a final memory usage of 915.49 MB. Here, peak memory refers to the maximum GPU memory allocated at any point during inference, while final memory indicates the memory remaining after inference. These results demonstrate that our method is both memory-efficient and scalable to datasets with thousands of genes.

**Figure 4:**
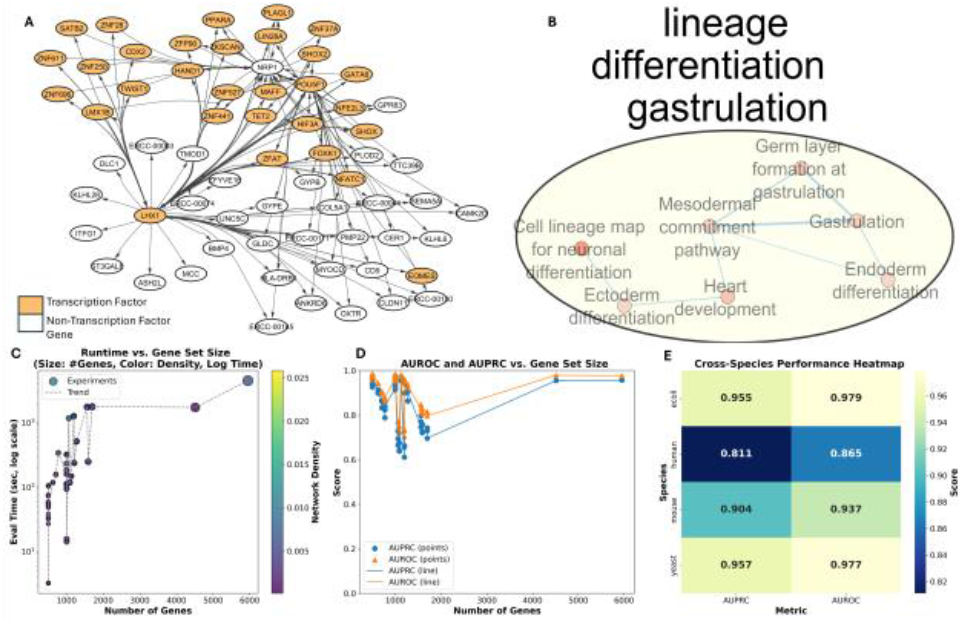
**(A)** novel regulatory interactions of hESCs predicted by GRNFormer outside of ground truth networks. **(B)** Enrichment map of GRN shown in 4A. **(C)** Runtime vs. number of genes. **(D)** Sampled_AUROC or Sampled_AUPRC vs. number of genes. **(E)** The average performance on the test datasets of four different species.

The prediction performance remained consistently high across increasing gene set sizes. Figure 4D shows Sampled_AUROC and Sampled_AUPRC scores as a function of the number of genes per dataset. GRNFormer maintained robust performance-even on large-scale networks-with both Sampled_AUROC and Sampled_AUPRC exceeding 0.9 for most settings beyond 2,000 genes. These results demonstrate that the model does not over-fit to low-dimensional settings and effectively captures regulatory signals in complex transcriptomic profiles.

To further assess generalizability across phylogenetic and regulatory variation, we benchmarked GRNFormer on gene expression datasets from *Homo sapiens, Mus musculus, Saccharomyces cerevisiae*, and *Escherichia coli*. As shown in Figure 4E, GRNFormer achieved high Sampled_AUROC and Sampled_AUPRC scores across all four species. This cross-species robustness highlights GRNFormer’s capacity to learn transferable regulatory representations. Together, these results affirm GRNFormer’s suitability for scalable and high-accuracy GRN inference-capable of generalizing across increasing gene dimensions, diverse species, and varied biological contexts without retraining or fine-tuning.

GRNFormer shows strong robustness to perturbations in gene expression data. Under 20 combinations of Gaussian noise and dropout, Sampled_AUROC and Sampled_AUPRC varied by less than 0.3% and 0.18%, respectively (Supplementary Results S4; Fig. S13; Tables S5.1–S5.3). This stability arises from correlation-preserving z-score normalization, TF-Walker’s context-based subgraph sampling, and the smoothing effect of GraViTAE’s variational latent space. These results demonstrate that GRNFormer maintains reliable predictive performance even under substantial noise and sparsity, supporting its applicability across diverse scRNA-seq conditions.

We further evaluated robustness of GRNFormer in comparisons to other benchmark existing SOTA methods, under the full-matrix setting (Sup Results S1.B). GRNFormer consistently maintained the highest median full AUROC and full AUPRC distributions across all datasets (Sup. Fig. S8), whereas competing methods exhibited substantially broader spread and lower lower-quartile performance. Average-rank and win-rate summaries (Sup. Fig. S9) confirm that GRNFormer achieves the greatest proportion of rank-#1 outcomes across datasets for both AUROC and AUPRC. In addition, paired t-tests confirm that GRNFormer significantly outperforms GNE and GNNLink across datasets (Supplementary Table S6).

### 3.6. Case study: GRNFormer Recovers hESC Pluripotency and Predicts PBMC Lineage Networks via Blind Inference

Human embryonic stem cells (hESCs) are pluripotent, self-renewing cells capable of differentiating into all somatic lineages (Gepstein 2002; Vazin and Freed 2010)(Gepstein 2002; Vazin and Freed 2010). Their shared regulatory programs with cancer-such as those controlling the cell cycle, apoptosis, and epigenetic states-make hESCs a valuable model for uncovering candidate oncogenic regulators (Blum and Benvenisty 2008)(Blum and Benvenisty 2008). In the first case study, to evaluate GRNFormer’s ability to recover biologically meaningful transcriptional circuits, we applied it to the hESC dataset that includes all annotated transcription factors and the top 500 most variable genes. The predicted GRN was bench-marked against cell-type-specific ChIP-seq ground truth, and all interactions shown in Supplementary Figure S14A correspond to predictions supported by the experimental ground truth reference. GRNFormer successfully reconstructed the core transcriptional architecture of hESCs, identifying master regulators of pluripotency, including *POU5F1 (OCT4), SOX2, MYC, and NANOG*, as central nodes in the network. These transcription factors are well established to operate in tightly interconnected autoregulatory circuits that coordinate gene expression programs essential for sustaining pluripotency and suppressing differentiation (Yeo and Ng 2013)(Yeo and Ng 2013). (See Supplementary Results S5, Supplementary Figure S14A-B for details)

In addition to predicting known pluripotency networks, GRNFormer revealed a novel transcriptional module centered on *GATA6, HAND1, NRP1, HNF1B*, and *TET2* (Figure 4A). These transcription factors are not typically active in ground-state pluripotency but are known to mediate early lineage specification events, including mesendoderm formation, cardiac development, neurulation, and vascular morphogenesis (Cirio et al. 2011; Lynch et al. 2025)(Cirio et al. 2011; Lynch et al. 2025). *TET2*, in particular, plays a role in DNA demethylation and chromatin remodeling, and its presence in this module suggests an epigenetically primed state (Eyres et al. 2021)(Eyres et al. 2021). Importantly, this subnetwork was absent from the considered ChIP-seq-derived hESC gold-standard networks, implying that GRNFormer can capture regulatory heterogeneity or transient cell states that elude bulk profiling techniques. To characterize the functional relevance of these predictions shown in Figure S14A and Figure 4A, we performed pathway enrichment analysis using g: Profiler(Raudvere et al. 2019)(Raudvere et al. 2019) and visualized the results ( Figure S14B and Figure 4B respectively) using Cytoscape(Shannon et al. 2003)(Shannon et al. 2003). The targets of the novel regulatory module (Figure 4B) were enriched for developmental programs, such as gastrulation and specification of the endoderm, mesoderm, and ectoderm (Muhr and Ackerman 2023)(Muhr and Ackerman 2023). Additional pathways included cardiac development and neuronal differentiation (Yao et al. 2017; Li et al. 2024; Mensah and Gowher 2024)(Yao et al. 2017; Li et al. 2024; Mensah and Gowher 2024). These findings indicate that the network in Figure 4A reflects an early, lineage-primed transcriptional program-potentially marking the transition from naïve pluripotency to germ layer commitment. The developmental programs enriched in the novel sub-network were absent from the core pluripotency GRN, suggesting that GRNFormer distinguishes stable from transitional regulatory states. These predictions reveal cryptic or low-frequency transcriptional programs in hESCs that may represent early cell fate determinants.

To illustrate interpretability, we examined a predicted subgraph centered on the gene NRP1 (Supplementary Results S6; Supplementary Figure S15). This NRP1-centered subgraph connects the gene to multiple transcription factors, *including POU5F1 (OCT4), LHX1, HAND1, GATA6*, and *TET2*. Higher predicted edge weights (shown in warmer colors) highlight strong model-inferred associations between NRP1 and both core pluripotency factors and lineage-priming TFs. Several high-weight edges are not present in the considered ChIP-derived gold standard but are consistent with literature links to developmental processes, suggesting that GRNFormer’s high-weight predictions are enriched for biologically plausible novel interactions. This case study highlights GRNFormer’s ability to recover both known and novel, biologically plausible regulatory modules.

In the second case study, to evaluate GRNFormer’s ability in resolving cell-type-resolved transcriptional programs, we applied it in a zero-shot setting to the 10x Genomics PBMC 3k dataset, a standard benchmark for immune single-cell profiling. The dataset was obtained in preprocessed form via the Scanpy toolkit, and GRNFormer inferred regulatory networks without any access to cell-type labels, clustering results, or pathway priors, relying solely on gene expression (Wolf, Angerer and Theis 2018)(Wolf, Angerer and Theis 2018).

A subnetwork extracted from the predicted GRN (Supplementary Figure S16A) highlights high-centrality genes with well-established immunological relevance, including *MS4A1, GNLY, NKG7, KLRB1, LST1, FCER1A*, and *TAL1*. These genes mark key immune lineages: *MS4A1 (CD20)* marks B cells (Stamenkovic and Seed 1988)(Stamenkovic and Seed 1988), *GNLY* and *NKG7* define cytotoxic *T* and *NK* cells(Krensky and Clayberger 2005)(Krensky and Clayberger 2005), *KLRB1* is linked to *NK* cell function(Fang and Zhou 2024)(Fang and Zhou 2024), *LST1* marks monocytes(Rollinger-Holzinger et al. 2000)(Rollinger-Holzinger et al. 2000), *FCER1A* labels plasmacytoid dendritic cells(Reshetnikova et al. 2018)(Reshetnikova et al. 2018), and *TAL1* regulates early hematopoietic lineage commitment(Real et al. 2012)(Real et al. 2012). These predictions underscores the model’s capacity to capture underlying regulatory logic directly from transcriptomic signals.

To interpret the functional landscape of the inferred network, we also performed pathway enrichment analysis using g:Profiler and visualized the results in Cytoscape via Enrichment Map plugin. The enriched pathways formed coherent modules aligning with known PBMC biology (Supplementary Figure S16B and Sup. Results S7),including vesicle trafficking, antigen presentation, and apoptotic signaling pathways. Interaction module around *TAL1* inferred by GRNFormer is supported by literature evidence, that *TAL1* is a master regulator of hematopoiesis (Sanda et al. 2012)(Sanda et al., 2012).These findings demonstrate that GRNFormer can infer biologically coherent cell-type-specific functional modules from single-cell data without external supervision. The model not only recapitulates known transcriptional circuits in immune lineages but also organizes them into interpretable regulatory programs, validating its utility for de novo GRN reconstruction in complex tissues.

## 4 Discussion

GRNFormer offers a scalable, context-aware framework for inferring gene regulatory networks (GRNs) from transcriptomic data. Unlike traditional approaches that rely on predefined motifs, GRNFormer learns regulatory patterns through TF-centered subgraphs sampled via the TF-Walker strategy, enabling inductive generalization. Ablation studies confirmed the critical role of this approach, with random sampling resulting in diminished performance. The Gene Transcoder and GraViTAE modules further capture biologically meaningful features and learn gene regulatory representations without relying on sequence priors.

The model demonstrates effective generalization across the species evaluated (human, mouse, yeast, E. coli), data types (bulk and single-cell RNA-seq), and regulatory modalities. On DREAM5 yeast and bacterial benchmarks-despite no prior exposure to prokaryotic data-GRNFormer achieved high Sampled_AUROC (~0.97) and Sampled_AUPRC (~0.95)., full test-set AUROC/AUPRC and greater EPR win rate (~84%). It also uncovered lineage-specific regulatory programs in human tissues and identified functional modules linked to pluripotency, antigen presentation, Epithelial–Mesenchymal Transition (EMT), and developmental pathways such as gastrulation. Additionally, cross-context interactions, including links between pluripotency and oncogenesis, were also revealed. Beyond performance, GRNFormer provides practical advantages in data efficiency, scalability, and interpretability. Unlike many supervised GRN methods, it generalizes to new species without retraining in our evaluations, making it applicable to systems with limited annotation. Its modular design allows easy adaptation, while edge-centric decoding ensures computational scalability without full-graph propagation.

Nevertheless, GRNFormer has certain limitations. Although GRNFormer performs well in zero-shot and cross-context settings, its accuracy may be reduced in datasets with extreme sparsity or weak transcription factor signal. Moreover, its reliance on local expression neighborhoods may overlook distal or chromatin-level regulatory interactions such as enhancer– promoter loops.

Looking forward, future developments could extend GRNFormer’s capabilities in several directions. Integrating multi-omics data, such as chromatin accessibility (ATAC-seq), epigenetic modifications, and TF motif occupancy, could enrich the contextual learning of regulatory interactions. Similarly, adapting the architecture to model temporal dynamics would enable causal modeling of transcriptional cascades. Further, incorporating cross-cell alignment or contrastive learning objectives may improve generalization in datasets with limited signal or across species.

## Acknowledgements

This work was supported by NSF grants (#: CCF2343612 and IOS2525780) and a Department of Energy grant (#: DE-SC0026121).

