## supplementary document for "GRNFormer: Accurate Gene Regulatory Network Inference Using Graph Transformer"

### Supplementary Methods and Figures

#### S1. GCEN Construction

A detailed description of GCEN generation appears in the main manuscript (Methods, Section *GCEN Construction*). Supplementary Fig. S1A summarizes the workflow, including correlation computation, thresholding, and graph assembly.

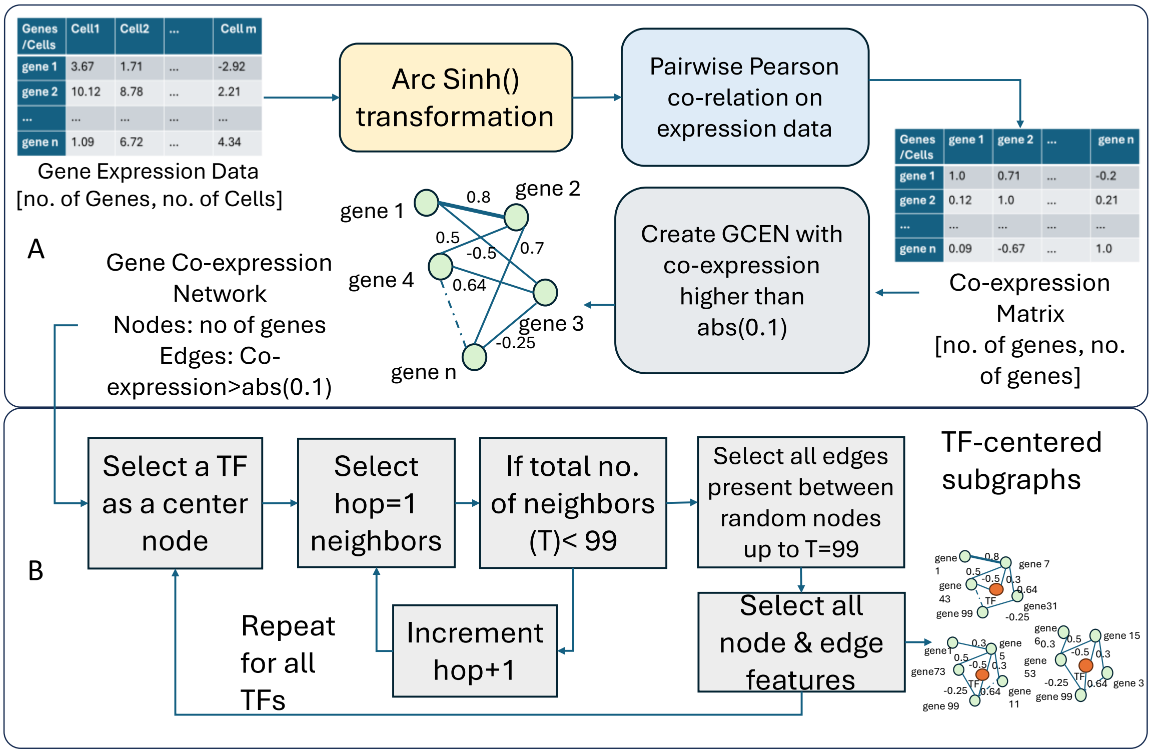


Figure S1. Input data preparation. (A) Gene expression preprocessing steps and gene co-expression network generation; (B) TF-walker methodology used for subgraph sampling during training of GRNFormer.

#### S2. TF-Walker (Training-Time) Sampling Formalization

The TF-Walker sampling algorithm is shown in Supplementary Fig. S1B and formalized as follows:

Let $G = (V, E)$ be the GCEN, $\mathrm{dist}_G(i, j)$ the shortest-path distance between genes iand jin G, and $S_{max}$ (shows as T in Fig.S1B) the maximum subgraph size.

For each transcription factor v, the h-hop neighborhood is defined as

$$\mathcal{N}_h(v) = \{ i \in V : \mathrm{dist}_G(v, i) \leq h \}.$$

TF-Walker increases the hop radius h until

$$| \mathcal{N}_h(v) | \geq S_{max} \text{or} \mathcal{N}_h(v) = \mathcal{N}_{h-1}(v).$$

If $| \mathcal{N}_h(v) | \leq S_{max} - 1$, all genes in the neighborhood are retained. If $| \mathcal{N}_h(v) | > S_{max} - 1$, we uniformly sample a subset without replacement

$$\mathcal{S}_v \subset \mathcal{N}_h(v), | \mathcal{S}_v | = S_{max} - 1,$$

and construct the induced TF-centered subgraph with node set and edge set

$$V_v = \{v\} \cup \mathcal{S}_v, E_v = \{ (i, j) \in E : i, j \in V_v \}.$$

Neighborhood extraction and hop-wise expansion are implemented using the k_hop_subgraph operator in PyTorch Geometric, which ensures efficient computation of shortest-path neighborhoods and consistent mapping of node indices during subgraph induction.

#### S3. Gene-Transcoder Algorithm

The algorithm of the Gene-Transcoder, described in the main manuscript (Methods, Gene-Transcoder), is shown in Supplementary Fig. S2.

Notation

- $B$: batch size; $m$: number of genes in each TF-centered subgraph (here $m = 100$); $T$: number of cells
- $d_{\text{model}}$: embedding dimension (e.g. 64); $H$: number of attention heads (e.g. 4)
- $X_{\text{in}} \in \mathbb{R}^{B \times m \times (T+1)}$: input tensor where, for each batch $b$ and gene $i$, $X_{\text{in}}(b, i, :) = (x_{b,i,1}, \dots, x_{b,i,T+1})$ containing expression of gene $i$ across $T$ cells and a TF-identity flag
- $X \in \mathbb{R}^{B \times m \times d_{\text{model}}}$: output tensor of fixed-size gene embeddings
- $W_{\text{conv}}, b_{\text{conv}}$: parameters of the 1D convolution; $W_h^Q, W_h^K, W_h^V, W^O$: multi-head attention projection matrices; $W_1, W_2, b_1, b_2$: feed-forward network parameters

**Algorithm: Gene-Transcoder**
**Input:** Gene expression matrix $X_{\text{in}} \in \mathbb{R}^{B \times m \times (T+1)}$.
For each batch index $b$ and gene index $i$, $X_{\text{in}}(b, i, :) = (x_{b,i,1}, \dots, x_{b,i,T+1})$ contains the gene expression across cells and the TF-identity flag.
**Output:** Fixed-size gene embeddings $X \in \mathbb{R}^{B \times m \times d_{\text{model}}}$.

**1. 1D Convolution Along the Cell Axis**
For each batch $b$, gene $i$, position $t \in \{1, \dots, T + 1 - K + 1\}$ and channel $c \in \{1, \dots, d_{\text{model}}\}$, apply a temporal convolution:

$$h_{b,i,t,c}^{(0)} = \sum_{k=0}^{K-1} w_c\,(k)\, x_{b,i,t+k} + b_c.$$

Stacking over time $t$ and channels $c$ gives:

$$H_{b,i}^{(0)} = \left(h_{b,i,1,:}^{(0)}, \dots, h_{b,i,T+1,:}^{(0)}\right) \in \mathbb{R}^{(T+1) \times d_{\text{model}}}.$$

Collecting all batches and genes: $H^{(0)} \in \mathbb{R}^{B \times m \times (T+1) \times d_{\text{model}}}$. In implementation $K = 1$, so the convolution reduces to a per-cell linear projection.

**2. Transformer Encoder**
For each $(b, i)$ treat $H_{b,i}^{(0)} \in \mathbb{R}^{(T+1) \times d_{\text{model}}}$ as a sequence of length $(T + 1)$.
**Multi-head self-attention:** For each head $h = 1, \dots, H$:

$$Q_h = H_{b,i}^{(0)} W_h^Q, \quad K_h = H_{b,i}^{(0)} W_h^K, \quad V_h = H_{b,i}^{(0)} W_h^V,$$

with $W_h^Q, W_h^K, W_h^V \in \mathbb{R}^{d_{\text{model}} \times d_k}$. Attention output:

$$A_h = \text{softmax}\left(\frac{Q_h K_h^\top}{\sqrt{d_k}}\right) V_h.$$

Concatenation and projection:

$$\widetilde{H}_{b,i} = \text{Concat}(A_1, \dots, A_H)\, W^O,$$

where $W^O \in \mathbb{R}^{(Hd_k) \times d_{\text{model}}}$.
Position-wise feed-forward network:

$$H_{b,i}^{(1)} = \left(\max(0, \widetilde{H}_{b,i} W_1 + b_1)\right) W_2 + b_2,$$

where $W_1 \in \mathbb{R}^{d_{\text{model}} \times d_{\text{ff}}}$, $W_2 \in \mathbb{R}^{d_{\text{ff}} \times d_{\text{model}}}$. Overall: $H^{(1)} \in \mathbb{R}^{B \times m \times (T+1) \times d_{\text{model}}}$.

**3. Mean Pooling Across Cells**

$$x_{b,i,:} = \frac{1}{T+1} \sum_{t=1}^{T+1} H_{b,i,t,:}^{(1)}, \qquad x_{b,i,:} \in \mathbb{R}^{d_{\text{model}}}.$$

Stacking over all $b$ and $i$ yields final embeddings: $X \in \mathbb{R}^{B \times m \times d_{\text{model}}}$.
**Output:** Return $X$, the fixed-size gene embeddings used as inputs to the GraViTAE module.

Figure S2: Algorithm of Gene-Transcoder

#### S4. TransConv and Pairwise Attention

The TransConv block is shown in Supplementary Fig. S3A, and the pairwise attention, message passing, and update step implemented through the graph transformer convolution layer are shown in Supplementary Fig. S3B.

For each gene as a node, attention weights are computed by comparing the query representation of a target gene with key representations of its neighbors, modulated by their connecting edge features. This pairwise attention mechanism allows the model to selectively weigh incoming messages based not only on the features of neighboring genes but also on the biological plausibility of their interactions. The resulting attention scores guide message passing, enabling the model to distinguish between biologically meaningful and spurious links.

As shown in Figure S3B, The pairwise attention mechanism computes the importance of a neighboring node ( j ) a nd its connecting edge ($e_{ij}$ ) relative to a target node ( i ). For the target node ( i ), a Query Vector (Q) is computed by transforming its feature vector ($X_i$) using a learnable weight matrix ($W_3$):

$$Q = W_3 X_i.$$

Similarly, a Key Vector (K) for a neighboring node (j) is calculated as:

$$K = W_4 X_j.$$

To explicitly incorporate edge information, the edge feature ($E_{ij}$), representing the relationship between nodes (i) and (j) is transformed using a weight matrix ($W_6$ ). The combined representation of the key vector (K) and edge feature ($E_{ij}$) is expressed as:

$$K + W_6 E_{ij}.$$

The attention mechanism then computes the attention score ($a_{ij}$),representing the relative importance of node (j) and its connecting edge ($e_{ij}$) to node (i). This is done using a scaled dot-product operation:

$$a_{ij} = \text{softmax}\left(\frac{Q \cdot (K + W_6 E_{ij})}{\sqrt{d}}\right),$$

where (d ) is the dimensionality of the feature space and serves as a scaling factor to stabilize the computation of dot products. The pairwise attention mechanism ensures that both node and edge features contribute meaningfully to the attention computation, allowing the model to learn the nuanced importance of edges within the graph.

Message Passing and Node Feature Update

Once the attention scores ($a_{ij}$) are computed, they are used to aggregate information from neighboring nodes. Each neighboring node (j) sends a message to the target node (i). This message is constructed using the Value Vector (V), which is derived by transforming ($X_j$) as:

$$V = W_2 X_j.$$

The message is further enriched by the transformed edge feature ($W_6 E_{ij}$), resulting in the final message term:

$$\alpha_{ij} = a_{ij} \cdot \left(V + W_6 E_{ij}\right),$$

where the attention score ($a_{ij}$) adaptively scales the contribution of the message from (j) based on the pairwise importance between ( j ),( i ), and their connecting edge ($e_{ij}$)

The updated node feature ($X_i$ ') is obtained by combining the original feature ($W_1 X_i$) with the aggregated messages from all its neighbors:

$$X_i' = W_1 X_i + \sum_{j \in \mathcal{N}(i)} \alpha_{ij}$$

where (N(i)) is the set of neighbors of node (i). This formulation ensures that the updated feature of node (i) is informed not only by its own attributes but also by the weighted contributions from its local neighborhood, incorporating both node-to-node and edge-based relational information. Similarly, the edge features are updated independently as:

$$E_{ij}' = W_6 E_{ij}.$$

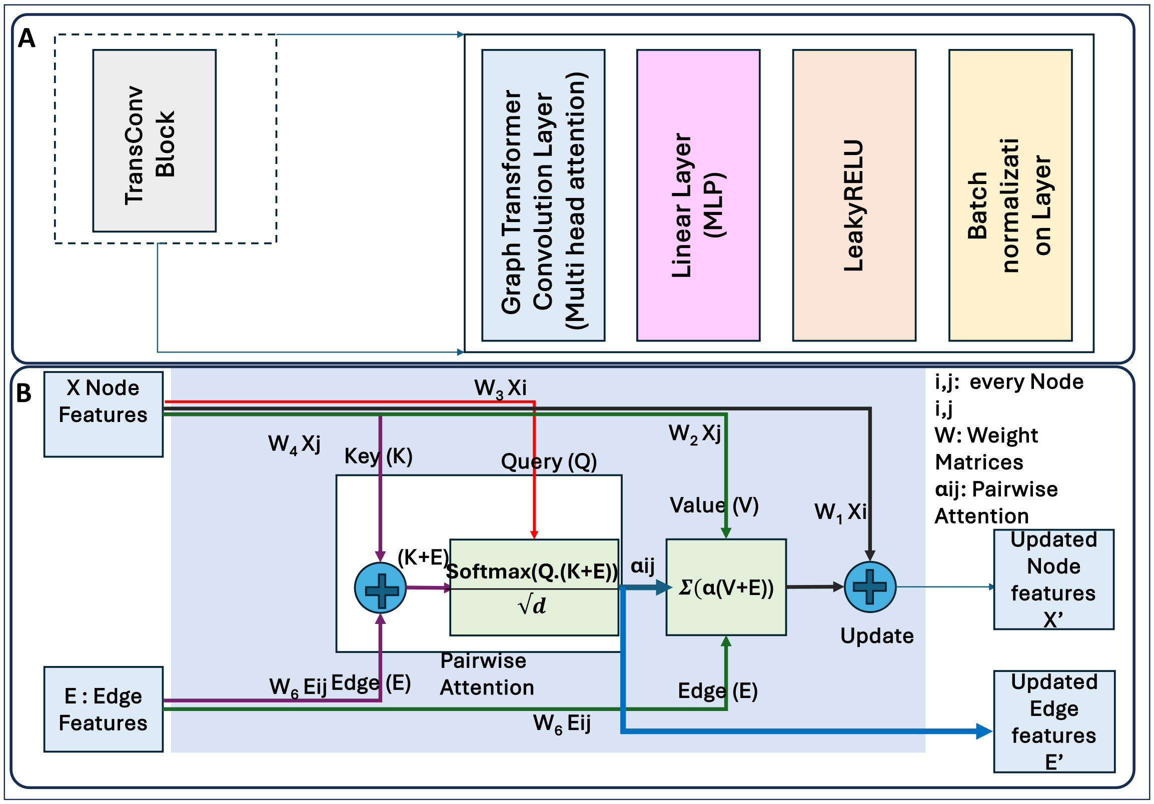


Figure S3: A) Detailed architecture of TransConv Block B) Block representation of pairwise attention used in Transformer Convolution layer

#### S5. Adjacency Matrix Reconstruction

The decoder outputs two components: node embeddings $Z' \in \mathbb{R}^{(Nxd)}$ , where $N$ stands for number of genes in the subgraph, encoding each gene's regulatory identity, and edge representations $E' \in \mathbb{R}^{(2xMxd)}$ where $M$ stands for number of edges present in input subgraph, capturing pairwise regulatory features. $d$ is the dimension of decoder output. To estimate regulatory interactions, an initial adjacency matrix $A_0$ is computed via the inner product of node embeddings:

$$A_0 = Z'(Z')^T$$

This operation yields a symmetric interaction matrix, where each element $A_0[i,j]$ reflects the interaction strength between gene $i$ and gene $j$, as inferred from their respective representations. While regulators and targets may occupy different contexts, genes that are likely to interact are embedded such that their inner products are higher. This approach aligns with established graph representation learning frameworks, such as Variational Graph Auto-Encoders (Kipf and Welling, 2016).

To enrich these pairwise estimates with contextual information captured at the edge level, the model applies mean pooling across the edge dimension of E′ and integrates the resulting values into the corresponding positions of the adjacency matrix. Let $\bar{E} \in \mathbb{R}^{NxN}$ denote this pooled edge tensor, where each entry represents an aggregated edge feature score between gene pairs. The combined interaction matrix becomes:

$$A = A_0 + \bar{E}$$

This combination allows the model to refine predictions for regulator-target pairs with distinct embeddings. Although the node-based inner product is symmetric, the addition of edge-specific embeddings introduces asymmetry, enabling directed regulatory interactions. Finally, a sigmoid activation function is applied elementwise to the combined matrix to transform the raw interaction scores into probabilistic values bounded between 0 and 1:

$$P = \text{Sigmoid}(A),$$

where $P[i,j] \in [0,1]$ indicates the predicted probability of a regulatory relationship between gene *i* and gene *j*.

#### S6. Variational Formulation & Loss Functions

**Variational formulation.**

For each transcription factor v, TF-Walker extracts a TF-centered subgraph with

$x^{(v)}$:node and edge features of the subgraph, $y^{(v)}$:TF–target labels for gene pairs $(i, j)$, $z^{(v)}$:latent representation of the subgraph.

The encoder defines a Gaussian posterior

$$q_\phi(z^{(v)} \mid x^{(v)}) = \mathcal{N}(z^{(v)} \mid \mu_\phi(x^{(v)}), \sigma_\phi(x^{(v)})^2 I),$$

where $\mu_\phi(x^{(v)})$and $\sigma_\phi(x^{(v)})$are the encoder-predicted mean and standard deviation vectors, and Iis the identity matrix.

Sampling uses the standard reparameterization rule,

$$Z^{(v)} = \mu_\phi(x^{(v)}) + \sigma_\phi(x^{(v)}) \odot \epsilon, \epsilon \sim \mathcal{N}(0, I),$$

where $\odot$denotes element-wise multiplication and $Z^{(v)}$ stands for latent embedding sampled for the TF-centered subgraph.

The evidence lower bound (ELBO) for TF vis

$$\mathcal{L}_{\text{ELBO}}^{(v)} = \mathbb{E}_{q_\phi(z^{(v)} \mid x^{(v)})} [\log p_\theta (R^{(v)} \mid z^{(v)})] - \text{KL} (q_\phi(z^{(v)} \mid x^{(v)}) \parallel \mathcal{N}(0, I)),$$

**where** $p_\theta(R^{(v)} \mid z^{(v)})$ is the decoder, and $R^{(v)}$ denotes the reconstructed node and edge representations produced by the decoder, which are subsequently transformed by the edge-reconstruction module to obtain TF–target probabilities $\hat{y}_{ij}^{(v)} \in [0,1]$for each gene pair $(i, j)$in the TF-centered subgraph.

**Loss Functions**

The binary cross-entropy loss evaluates the discrepancy between the predicted edge probabilities $\hat{y}_{ij}$ and the true binary labels $y_{ij} \in \{0,1\}$ for each gene pair (i,j) , and is computed as:

$$\mathcal{L}_{\mathrm{BCE}} = -\sum_{(i,j)} \left[ y_{ij} \cdot \log\left(\widehat{y_{ij}}\right) + \left(1 - y_{ij}\right) \cdot \log\left(1 - \widehat{y_{ij}}\right) \right]$$

To enforce regularization in the variational latent space, we incorporate the Kullback–Leibler divergence between the approximate posterior q(z|x) (parameterized by the encoder) and a standard normal prior p(z). This is given by:

$$\mathcal{L}_{\mathrm{KL}} = -\frac{1}{2} \sum_{i=1}^{n} \left(1 + \log(\sigma_i^2) - \mu_i^2 - \sigma_i^2\right)$$

Where $\mu_i$ and $\sigma_i^2$ are the mean and variance of the latent embedding for gene i.

The total loss function is the weighted sum of these two terms:

$$\mathcal{L}_{total} = \mathcal{L}_{BCE} + \beta \cdot \mathcal{L}_{\mathrm{KL}}$$

where β is a scaling factor with value 0.01 or 1/(#genes in subgraph) controlling the strength of KL regularization.

#### S7. GRN Inference Strategy – Deterministic TF-Walker Expansion Algorithm

1. **Initialize**
   - Input: Graph G = (V, E), transcription factors (TFs) as a list, target subgraph size T = 99.
   - Output: Subgraphs S1, S2, ..., Sk.

2. **For each TF v ∈ TF list:**
   **Step 2.1: Identify 1-hop neighbors.**
   - Find all 1-hop neighbors N1(v) of v, v is a TF node, and N1(v) is a 1-hop neighbors of v.

   **Step 2.2: Check the size of N1(v):**

   - **Case 1: |N1(v)| < T:**
     - Increment hops by 1.
     - Include neighbors from hop + 1 until |N(v)| ≥ T (where N(v) is the set of unique neighbors).
     - Proceed to continue with subgraph creation using N(v).
   - **Case 2: |N1(v)| ≥ T**:
     - **Subcase 2.1**: N1(v) is too large to fit into a single subgraph:
       - Divide N1(v) into multiple subgraphs S1, S2, ..., such that each subgraph contains ≤ T nodes.
       - If any subgraph Si has fewer than T nodes, add nodes from hop + 1 neighbors to reach T.
     - **Subcase 2.2**: If a single subgraph is to be created with a mix of neighbors from different depths:
       - Assign the first T/2 nodes to S1 from 1-hop neighbors.
       - Assign the remaining T/2 nodes from hop + 1 neighbors to S1.
       - If N1(v) and Nk(v) (incrementally) still do not meet the target size T:
         - Increment the hop and include additional k-hop neighbors iteratively until T-sized subgraphs are satisfied.
         - For each depth, maintain balanced selection by ensuring T/2 allocation from N1(v) and subsequent depths.

3. **Construct subgraphs for each TF v:**
   - Include all edges Es ⊆ E where both endpoints belong to the subgraph nodes.
   - Include all node features and edge features for the selected nodes and edges.

4. **Process remaining uncovered nodes in V:**
   - Identify nodes Vr that are not included in any subgraph.
   - Select |Vr| as center nodes and repeat the process until all nodes are covered.
   - For subgraphs with fewer than T nodes, increment the hop or randomly add nodes to meet the target size T.

5. **Output the final subgraphs**:

Subgraphs S1, S2, ..., Sk, where each subgraph has exactly T nodes (if possible), and all edges and features within the subgraphs are included.

Figure S4. Algorithm of real-time subgraph sampling for GRN inference.

#### S8. Evaluation metrics

AUROC quantifies the model's ability to distinguish regulatory from non-regulatory interactions across all thresholds. AUPRC emphasizes the model's accuracy in detecting true regulatory edges, making it especially informative for imbalanced datasets.

F1 score balances precision and recall, offering a single summary of model performance when both false positives and false negatives matter. Precision reflects the fraction of predicted regulatory edges that are correct, while recall captures the proportion of true regulatory interactions successfully recovered. F1 score, precision, and recall depend on the decision threshold used to select positive predictions. The threshold leading to the maximum F1 score is used. Although accuracy (percentage of correct predictions in all predictions) is also reported, it is less reliable in imbalanced settings and used primarily for completeness. Early Precision (EP) evaluates how accurately a GRN inference method prioritizes true regulatory interactions among the highest-confidence predicted edges. This metric focuses on the top-ranked predictions that are most biologically actionable, complementing global measures such as AUROC and AUPRC.

#### S9. Benchmark Setup

For comparative benchmarking, we evaluated GRNFormer against a representative set of classical and deep learning-based GRN inference methods. To ensure reproducibility and ease of deployment, some of the methods were run using Dockized environments provided by BEELINE, where available.

Among classical approaches, we selected four widely used and computationally efficient methods: PIDC, PPCOR, LEAP, and SCENIC-RITES. These were chosen for their speed and scalability, as they completed inference on 500-gene inputs in under one hour. More computationally intensive methods (e.g., GENIE3(Huynh-Thu *et al.,* 2010), GRNVBM (Sanchez-Castillo *et al.,* 2018)) were excluded due to runtime limitations in large-scale evaluations.

Five deep learning state-of-the-art (SOTA) methods (GNE, GNNLink, CNNC, STGRNs, scGREAT) were reproduced using their recommended within-dataset training procedures and default hyperparameters. For each test cell type, we used the same train–validation–test splits across all competing methods to maintain consistency.

GRNFormer was trained once on a dataset spanning five cell types (hESC, hHep, mDC, mHSC-E, mHSC-GM) and evaluated without fine-tuning on two additional cell types (mESC, mHSC-L).

All methods were evaluated on the same held-out test positives using **bootstrapped negatives** sampled from the **clean negative evaluation pool** (see Supplementary Method S10 for construction details). Each method was assessed using **100 iterations of 1:1 bootstrapped sampling**, and metrics derived from this balanced protocol are reported as **Sampled AUROC** and **Sampled AUPRC**. Metrics derived from Full-matrix like evaluations are reported as Full test-set AUROC/AUPRC.

#### S10. Clean Negative Evaluation Pool Construction and Sampled and Full Metrics Evaluation Protocol

To provide a rigorous and unbiased comparison of GRNFormer with all state-of-the-art (SOTA) methods, we implemented a unified full-matrix evaluation protocol designed to eliminate sampling bias, prevent information leakage. This section describes the evaluation framework.

Unlike 1:1 sampled negative evaluation-which may inflate Sampled_AUROC/Sampled_AUPRC and is sensitive to sampling, variance-full-matrix evaluation assesses each model across the complete gene x gene edge space. i.e. all possible gene-gene pairs are considered in the evaluation of metrics. Since many competing methods need data-specific training, considering all possible edges may result in data leakage. So, for robust and fair evaluation, for every dataset, we first split all known TF–target positive edges into training, validation, and testing sets following a 70/10/20 ratio using fixed seeds. During training and validation, each model records the specific negative edges it uses for optimization (including all deep learning SOTA methods: CNNC, scGREAT, STGRNS, GNNLink, GNE).

To avoid reusing any negative edges seen during training or validation, we construct a **clean negative evaluation pool** defined as:

$$N_{\text{clean}} = \{(i,j) \text{ for all gene–gene pairs}\} - \{\text{train/val/test positives, train/val negatives, self-loops}\}.$$

This clean negative evaluation pool is used for both bootstrapping negative samples in calculation of sampled metrics, as well as full matrix evaluation.

For sampled evaluation, we sampled negative edges from the clean negative evaluation pool at a 1:1 positive-to-negative ratio via bootstrapping (100 draws) to obtain stable Sampled_AUROC/Sampled_AUPRC estimates.

Full test-set AUROC and full test-set AUPRC are computed on the union of the **test positives** and $N_{\text{clean}}$.

#### S11. Hyperparameter Sensitivity Analysis

We evaluated the sensitivity of TF-Walker's two effective hyperparameters-the GCEN correlation threshold and the TF-centered subgraph size-using three datasets of different scales (*E. coli*, mESC cell-type specific TF500, and mESC cell-type specific TF1000). These analyses quantify how GCEN sparsity and neighborhood size influence TF-Walker expansion behavior, GRN structural properties, computational requirements, and downstream GRNFormer performance.

##### A. GCEN Correlation Threshold Sensitivity.

We examined correlation thresholds of 0.1, 0.3, 0.5, and 0.7 using the mESC cell-type–specific TF500 dataset. Supplementary Figure **S5A-F** reports several structural and performance curves that together describe how thresholding alters GCEN topology.

The **threshold–true-edge coverage** curve (Supplementary Fig. S5A) measures the fraction of known TF–target edges preserved in the GCEN. Coverage decreases steeply beyond 0.3, showing that biologically supported interactions are progressively removed as the graph becomes sparser.

The **high-probability edges vs GCEN threshold** curve (Supplementary Fig. S5B) quantifies how many high-confidence GRNFormer predictions (probability $\geqslant 0.6$) remain present and connected within subgraphs as the co-expression threshold becomes stricter. This curve declines drastically immediately after 0.3, revealing early loss of model-relevant regulatory structure even before Sampled_AUROC/AUPRC begin to drop.

The **network density** curve (Supplementary Fig. S5C) summarizes global GCEN sparsity. Density decreases gradually between 0.1 and 0.3 but then falls sharply, consistent with widespread edge removal and fragmentation of co-expression modules.

Finally, **Sampled_AUROC and Sampled_AUPRC** (Supplementary Figs. S5D–E) are highest at a threshold of 0.1-0.3 and decrease monotonically at higher thresholds. These performance declines mirror the structural degradation observed in TF-degree, true-edge retention, high-probability edges, and network density.

The **TF-degree distribution** (Supplementary Fig. S5F), which reflects how many co-expression neighbors each TF retains, shows a sharp decline in TF connectivity at thresholds above 0.3. This indicates rapid neighborhood collapse and explains why TF-Walker more frequently terminates early under stricter thresholding.

Together, these analyses show that thresholds $\geqslant 0.3$ substantially reduce TF connectivity, biological edge content, global density, and model-relevant edges, leading to lower predictive accuracy. A threshold of **0.1-0.3** preserves the most complete and biologically meaningful GCEN and supports the most stable TF-Walker behavior. We adopt **0.1 as the default correlation threshold**, while allowing users to raise it during inference when stricter filtering is appropriate.

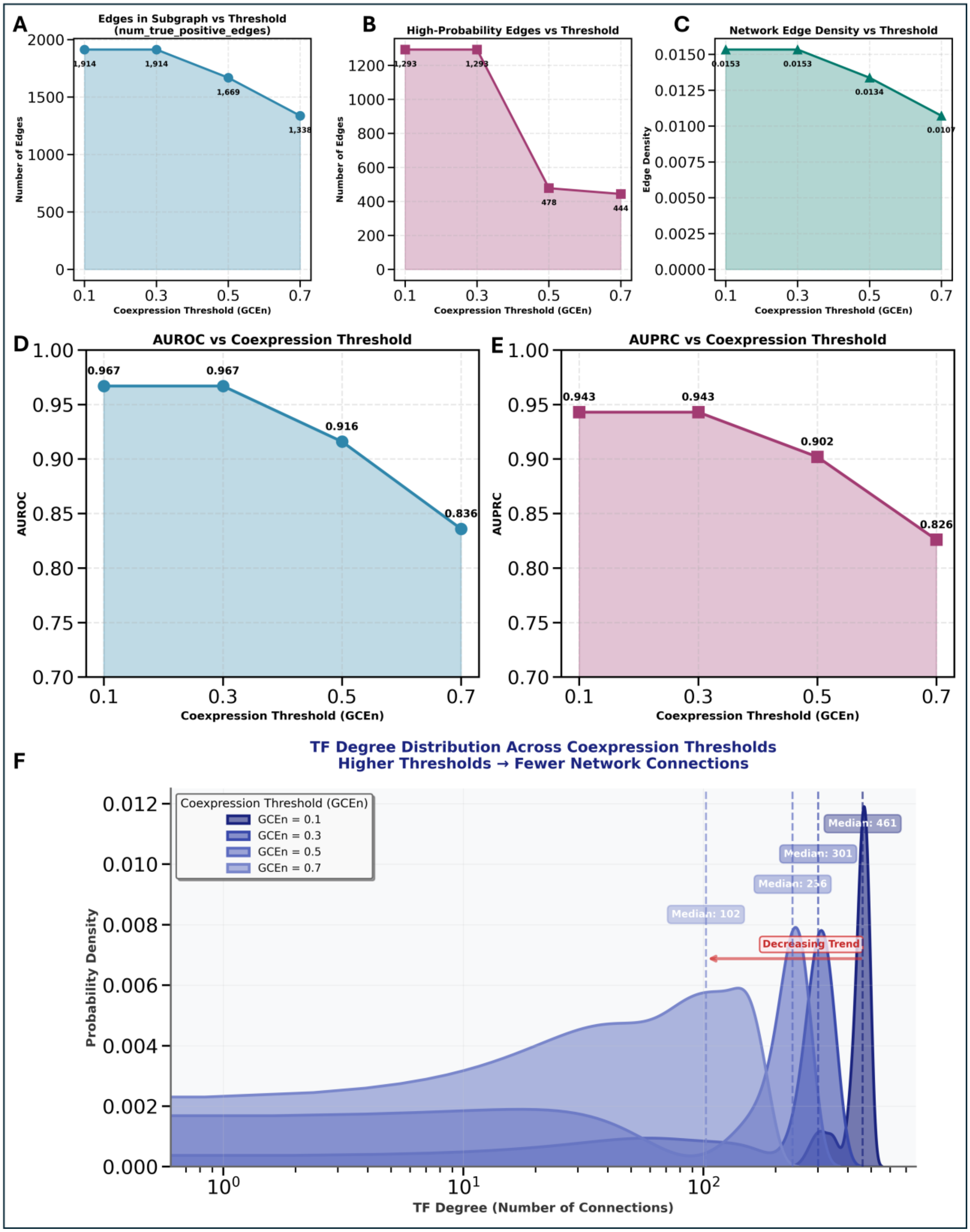


**Figure S5: Impact of co-expression thresholding on GCEN structure, TF connectivity, and GRNFormer performance (mESC TF500 dataset). (A)** *True positive regulatory edges retained vs. GCEN threshold.* Shows the number of experimentally supported TF–target edges preserved under each correlation cutoff. A steep decline at thresholds ⩾0.3 indicates loss of biologically meaningful regulatory relationships. **(B)** *High-probability edges retained vs. GCEN threshold.* Counts how many high-confidence GRNFormer-predicted edges remain present and connected within the subgraphs as the threshold increases. The abrupt reduction after 0.3 shows early removal of model-relevant edges before performance metrics decline.**(C)** *Global network edge density vs. GCEN threshold.* Depicts how overall GCEN sparsity changes with thresholding. Density decreases progressively from 0.1 to 0.3 and collapses at higher values, reflecting extensive edge loss and fragmentation of co-expression modules. **(D)** *AUROC vs. GCEN threshold.* GRNFormer achieves highest AUROC at threshold 0.1. Performance decreases as GCEN becomes sparser, consistent with the loss of TF neighborhood integrity. **(E)** *AUPRC vs. GCEN threshold.* AUPRC trends mirror AUROC: peak performance at threshold 0.1 followed by monotonic decline at higher thresholds. All AUROC and AUPRC are based on sampled evaluation **(F)** *TF-degree distributions across GCEN thresholds.* Probability density curves showing how TF connectivity shifts as thresholds increase. Median TF degree decreases from 461 (threshold 0.1) to <50 (threshold 0.7), illustrating rapid collapse of TF-centered neighborhoods.

#### B. Subgraph Size Sensitivity

We evaluated TF-Walker subgraph sizes of 10, 50, 100, 200, and 500 nodes using the *E. coli*, mESC cell-type–specific TF500, and mESC cell-type–specific TF1000 datasets to assess how neighborhood size affects representational quality and computational cost. Supplementary Figure **S6A–D** summarizes these effects.

The Sampled **AUROC and AUPRC** curves (Supplementary Figs. S6A–B) show that performance increases substantially from 10 to 100 nodes and remains stable between 100 and 200 nodes. Enlarging subgraphs to 500 nodes does not improve accuracy in any dataset. Across all conditions, GRNFormer maintains high performance (Sampled_AUROC > 0.96, Sampled_AUPRC > 0.93), indicating that the model is robust to the choice of subgraph size once sufficient TF–target context is included.

The **runtime** curve (Supplementary Fig. S6C) demonstrates that very small subgraphs require repeated TF-Walker expansions, increasing inference time. Larger subgraphs reduce the total number of required expansions and shorten runtime in dense networks such as mESC. In contrast, *E. coli* shows non-monotonic scaling due to variable neighborhood structures.

The **GPU memory usage** curve (Supplementary Fig. S6D) increases predictably with subgraph size. Memory demand remains moderate for 100–200 nodes but grows substantially at 500 nodes, reaching ~1.4 GB in E.coli. This pattern sets a practical upper limit on subgraph size for typical GPU hardware.

To summarize performance robustness quantitatively, **Supplementary Table S4** reports mean sampled AUROC/AUPRC, standard deviation, and coefficient of variation (CV) across all subgraph sizes and datasets. All datasets show very low variability (CV < 3%), and the Sampled_AUROC/AUPRC ranges are narrow, confirming that GRNFormer maintains stable accuracy across a 50-fold range in subgraph size. These statistics highlight the resilience of the model to neighborhood-size variation and reinforce the performance plateau observed between 100 and 200 nodes.

Together, the accuracy patterns, runtime behavior, memory scaling, and Table S4 statistics indicate that subgraph sizes in the **100–200 node range** offer the best trade-off between biological coverage, computational efficiency, and resource use. Subgraphs smaller than 50 nodes lack sufficient regulatory context and increase runtime, whereas very large subgraphs impose heavy memory costs without improving accuracy. Accordingly, we use **100 nodes as the default subgraph size** during training and recommend **100–200 nodes** for inference.

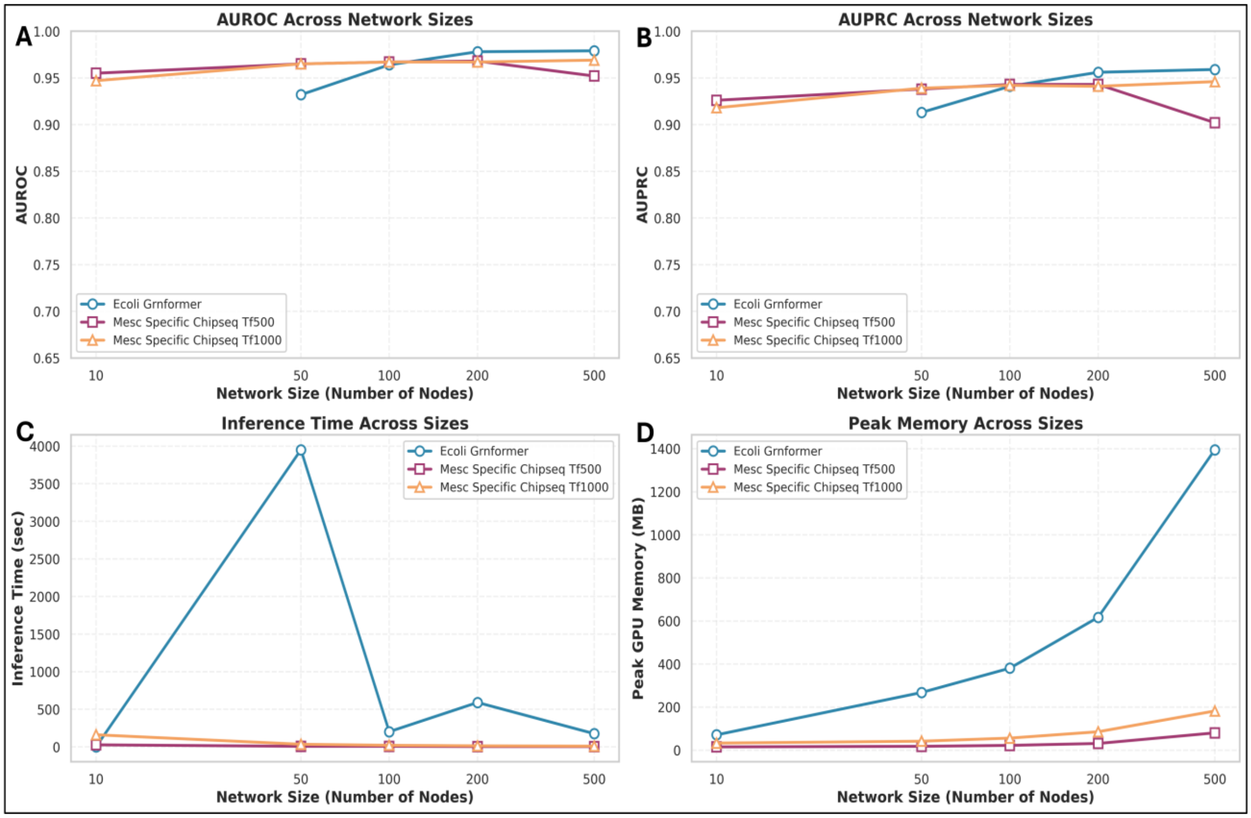


**Figure S6:** Effect of TF-Walker subgraph size on accuracy, inference time, and memory usage across three representative datasets (E. coli GRN, mESC cell-type–specific ChIP-seq TF500, and ChIP-seq TF1000). **(A)** *AUROC across subgraph sizes.* AUROC improves substantially from 10 to 100 nodes across all datasets and stabilizes between 100–200 nodes. No or a little additional performance gain is observed at 500 nodes. **(B)** *AUPRC across subgraph sizes.* AUPRC follows the same pattern as AUROC: strong improvement at small sizes, plateau at 100–200 nodes, and no benefit from very large subgraphs. All AUROC and AUPRC are based on sampled evaluation **(C)** *Inference time vs. subgraph size.* Inference becomes slower for very small subgraphs due to repeated TF-Walker expansions. Larger subgraphs reduce the total number of expansions and produce shorter runtimes in dense networks (mESC). The *E. coli* dataset displays non-monotonic behavior due to heterogeneous network connectivity. **(D)** *Peak GPU memory usage vs. subgraph size.* Memory usage increases steadily with subgraph size. Subgraphs of 100–200 nodes remain manageable for typical GPUs, while 500-node subgraphs require substantially more memory, reaching ~1.4 GB for mESC TF1000.

##### C. Recommended Parameter Settings

Based on these analyses, the following settings provide the most robust and efficient behavior across datasets:

- **GCEN correlation threshold:** Using **0.1** as the default. It preserves broad co-expression structure, retains true regulatory edges, and maintains high-probability predictions important for TF-Walker expansion.
- **TF-Walker subgraph size (training):** Use **100 nodes**, which provides reliable coverage and high accuracy with efficient memory use.
- **TF-Walker subgraph size (inference):** Use **100–200 nodes** depending on dataset size and hardware availability. Subgraphs <50 nodes risk incomplete neighborhoods, while subgraphs near 500 nodes substantially increase memory cost without benefits.

These recommendations reflect a balanced combination of biological fidelity, computational efficiency, and predictive stability across diverse GRN inference settings.

#### Supplementary Results and Figures

##### S1. Sampled & Full-Matrix Evaluation

##### A. Sampled Matrix Evaluation

To obtain unbiased performance metrics, we conducted 100 iterations of 1:1 bootstrapped negative sampling from the clean evaluation pool. In each iteration, every test positive edge was paired with a randomly drawn unseen negative edge. Because these metrics are derived from balanced sampling rather than the full edge space, they are explicitly reported as Sampled AUROC and Sampled AUPRC. This unified protocol eliminates sampling bias and ensures that all methods are compared under identical, strictly controlled evaluation conditions. In the sampled evaluation, test positives are restricted to TF-Walker–covered TF–target edges (i.e., gold-standard positives with at least one valid TF-Walker path). The accompanying software also reports the corresponding positive coverage, defined as the fraction of gold-standard positives that are TF-Walker–covered and included in the evaluation.

Supplementary Figure S7 presents heatmaps of Sampled AUROC (S7A) and Sampled AUPRC (S7B) across all methods and test datasets under this standardized bootstrapped evaluation. GRNFormer was evaluated on cell types that were not seen during training, whereas several baseline methods were trained and tested within the same cell type. Despite this, GRNFormer consistently achieves the highest or near-highest performance across nearly all datasets.

For Sampled AUROC (S7A), GRNFormer maintains strong performance in both non–cell-type-specific and cell-type-specific settings (Sampled_AUROC > 0.9), with particularly robust values observed in the mESC STRING benchmark and the ChIP-seq TF500/TF1000 gene sets. In terms of Sampled AUPRC (S7B), GRNFormer shows clear advantages in precision–recall behavior, especially on ChIP-seq datasets, where competing methods frequently exhibit substantial degradation.

Traditional statistical approaches such as PIDC, PPCOR, and SINCERITIES perform poorly under these settings, reflecting their limited ability to generalize across cell types. Among the competing methods, GNE demonstrates the strongest baseline performance, achieving the most competitive Sampled_AUROC and Sampled_AUPRC across datasets under the clean evaluation protocol. GNNLink performs well on several datasets for Sampled_AUROC, but its Sampled_AUPRC varies considerably, indicating sensitivity to dataset characteristics.

In contrast, GRNFormer surpasses all baselines—including GNE—on the majority of datasets, with particularly strong gains in Sampled AUPRC, where accurate identification of true regulatory relationships is most critical. These results indicate that GRNFormer reliably infers high-confidence gene regulatory networks (GRNs) in a data-agnostic manner and achieves state-of-the-art accuracy under a stringent evaluation framework in which all methods are tested against the same unseen positives and negatives.

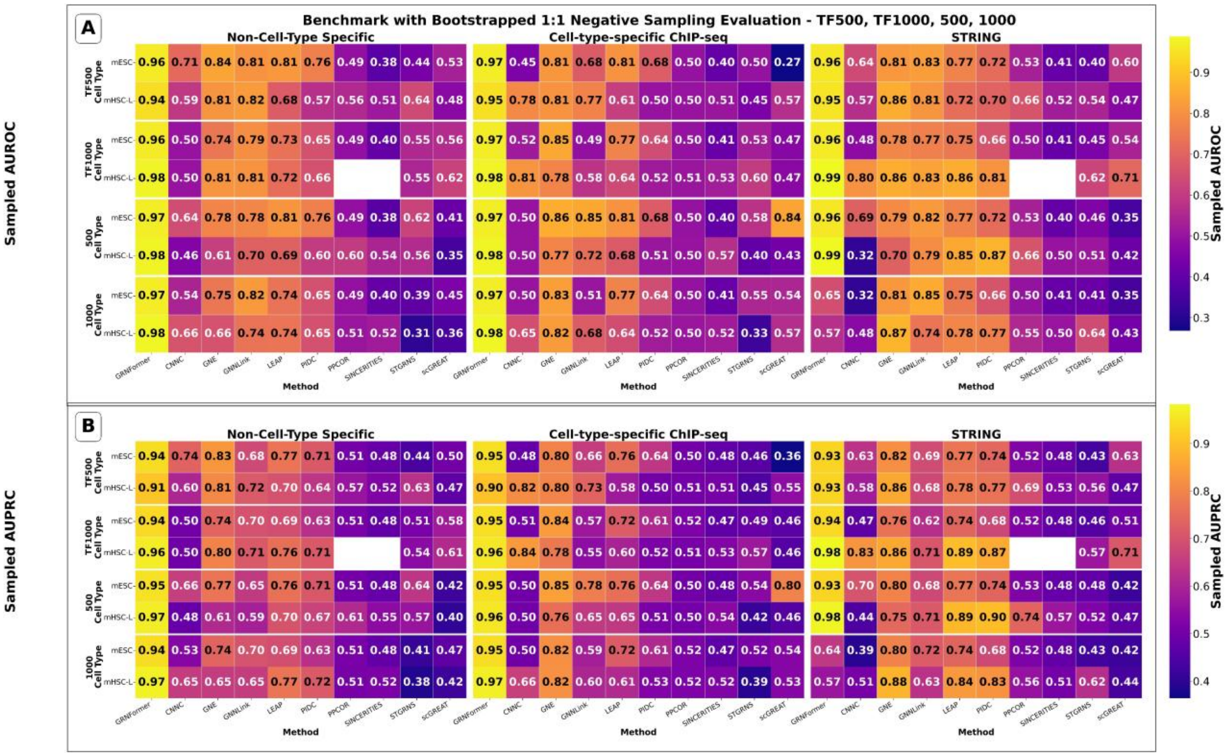


**Figure S7.** Sampled evaluation of GRNFormer and nine state-of-the-art methods across non–cell-type-specific, cell-type-specific ChIP-seq, and STRING datasets. (A) Sampled_AUROC computed over all test-set positive edges and the clean evaluation negative pool. (B) Sampled_AUPRC enabling comparison across datasets of different sparsity. GRNFormer achieves the highest or near-highest performance across nearly all datasets in both Sampled_AUROC /AUPRC, substantially outperforming existing state-of-the-art methods in cell-type-specific and non-cell-type-specific conditions.

#### B. Full-Matrix Evaluation

Following the full-matrix evaluation protocol described in Supplementary Method S10, we computed exhaustive AUROC, AUPRC, metrics using the clean negative evaluation pool. Supplementary Figure S8B illustrates the full test-set evaluation edge counts for all datasets to highlight test positives and clean evaluation negative pool. The resulting full test-set AUROC and AUPRC heatmaps are shown in Main manuscript Fig. 3A–B. Across more than half of the benchmark datasets, GRNFormer achieves the highest or near-highest performance under the most stringent evaluation setting.

We next examined the distributional behavior and robustness of these full-matrix metrics across heterogeneous biological contexts. Supplementary Fig. S8A displays the distributions of full test -set AUROC and full test-set AUPRC across all methods and datasets. GRNFormer attains the highest median values, exhibits the narrowest interquartile ranges, and shows markedly fewer low-performing outliers than competing approaches. These results indicate that GRNFormer performs strongly across most of the evaluated datasets.

To further quantify comparative performance, we assessed average-rank and win-rate statistics across all datasets, summarized in Supplementary Fig. S9. GRNFormer achieves the best overall average rank for both full test-set AUROC and full test-set AUPRC and obtains the highest win rate. This trend suggests that GRNFormer remains competitive across a broad range of evaluation scenarios.

Additionally, to formally assess whether the performance differences between GRNFormer and the strongest competing methods are statistically meaningful, we conducted paired **t-tests** across all benchmark datasets, comparing GRNFormer against **GNE** and **GNNLink**—the two competing methods that showed the closest performance under the evaluation pipeline. Tests were performed separately for (i) **bootstrap-based sampled AUROC/AUPRC**, and (ii) **full-matrix AUROC, AUPRC, and AUPRC-ratio**, using per-dataset paired differences for each metric. For additional interpretability, we also report **mean performance differences**, **win rates** (percentage of datasets where GRNFormer outperforms the competing method), and **p-value–based significance levels**.

The results, summarized in Supplementary Table S6, show that GRNFormer is significantly more accurate than both GNE and GNNLink across nearly all evaluation settings. Under the bootstrap sampled evaluation, GRNFormer achieves large positive mean differences, extremely high win rates (91.7%), and highly significant p-values ($p < 0.001$). Under full-matrix evaluation—which is more stringent—GRNFormer remains significantly superior, particularly relative to GNNLink, with strong mean AUROC and AUPRC-ratio advantages and high win rates (≥83%). Although differences relative to GNE are smaller in the full setting, GRNFormer still achieves statistically significant gains in AUROC ($p = 0.0283$) and consistent directional improvement across datasets.

These analyses confirm that GRNFormer's improvements over the strongest competing method are not only numerically large but also **statistically robust**, supporting the reliability of its performance advantages across diverse datasets and evaluation protocols.

Collectively, Supplementary Figs. S8–S9 and Supplementary Table S6 show that GRNFormer achieves high accuracy under full-matrix evaluation and demonstrates robust performance across the datasets analyzed relative to GNE, GNNLink, and other classical and neural benchmarking methods.

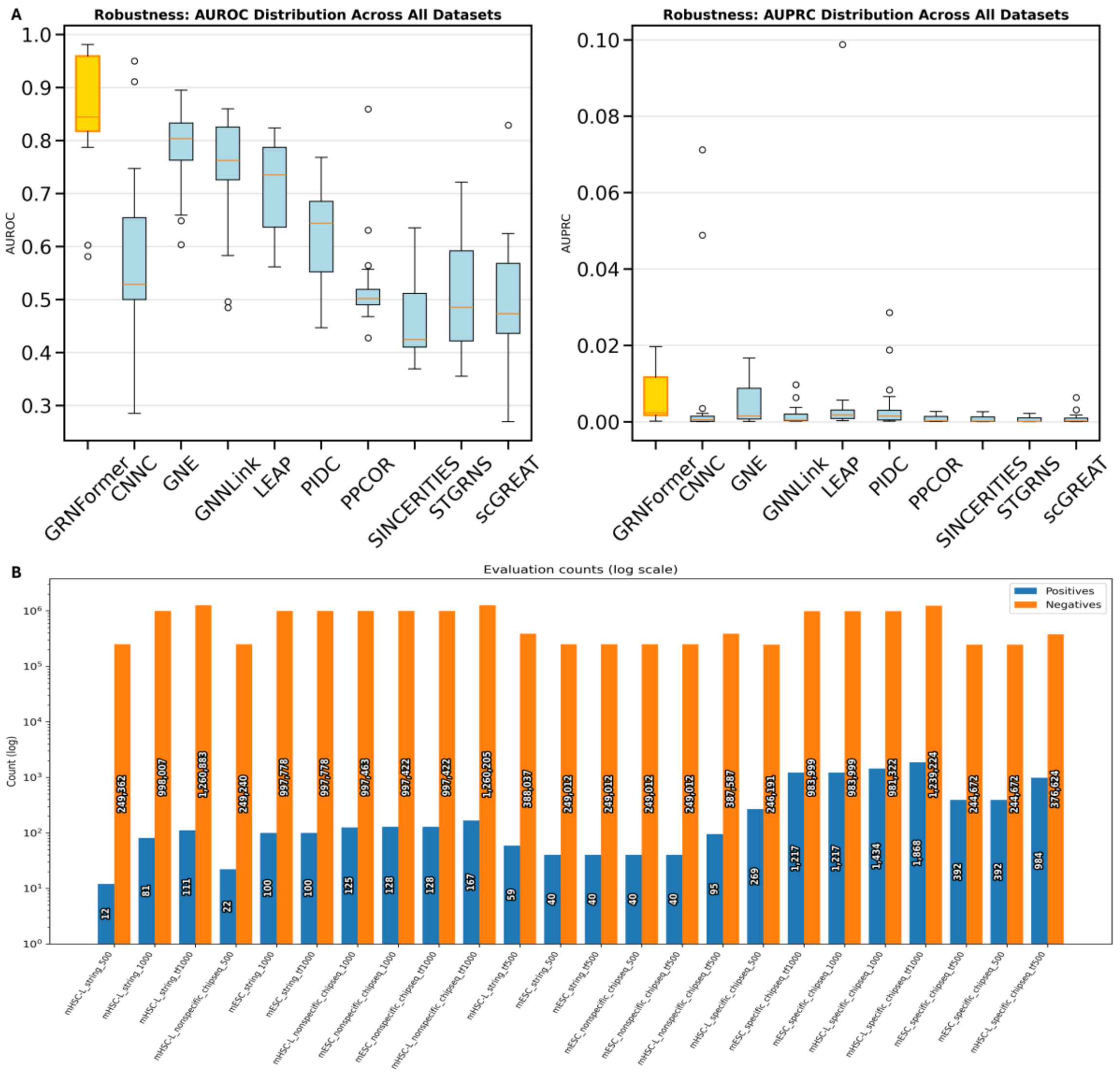


**Figure S8.** A. Distributional robustness of full-test-set AUROC and AUPRC across all datasets for GRNFormer and competing methods. Boxplots illustrate performance dispersion across 10 datasets. GRNFormer exhibits the highest median and the narrow interquartile range for both metrics, indicating consistently strong performance and low variability. Competing methods show substantially wider variance, with several methods exhibiting dataset-specific collapse. These figures demonstrate that GRNFormer's performance is not dataset-dependent but reflects a robust and generalizable modeling of transcriptional regulation. B. Bar plots to present the positive and negative edge counts of all the test datasets included for Full test-set AUROC/AUPRC calculations.

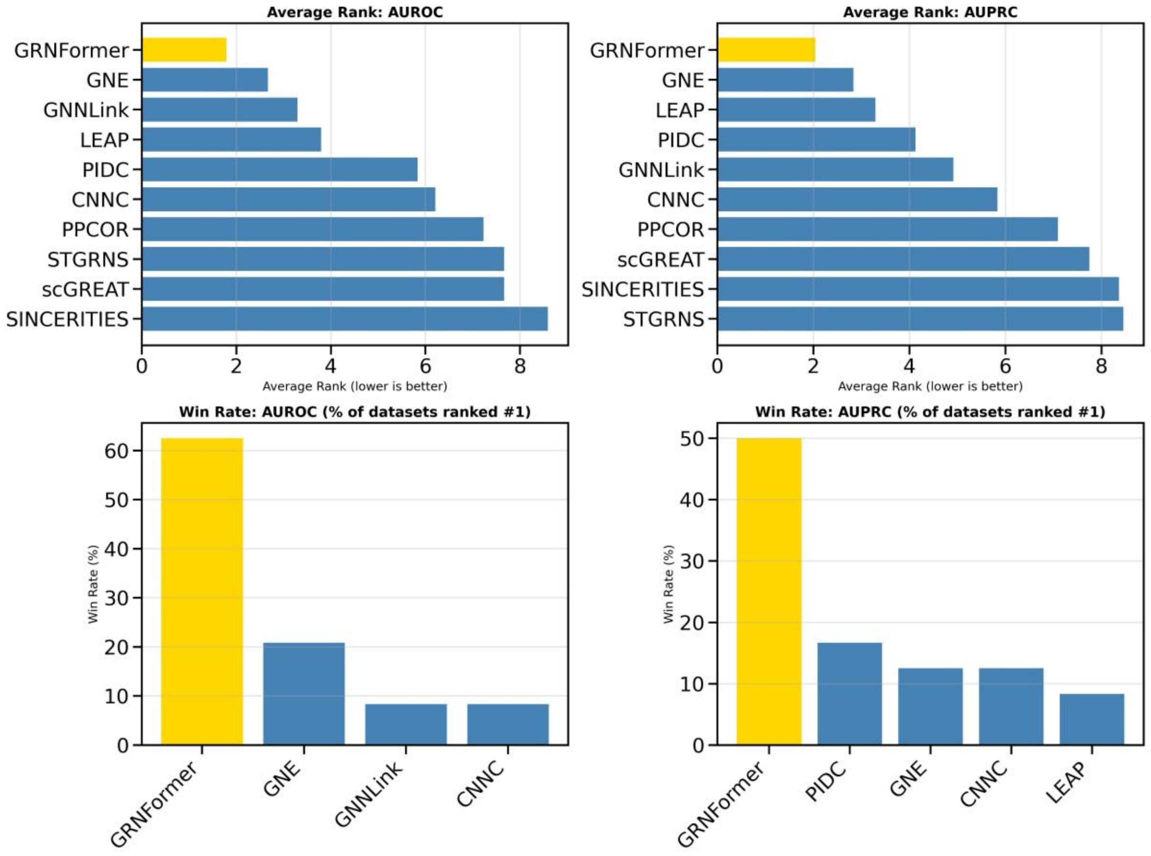


**Figure S9.** Summary ranking and win-rate analysis across all benchmark datasets. Top: average rank for AUROC (left) and AUPRC (right), where lower ranks indicate stronger performance. GRNFormer attains the best average rank across both metrics. Bottom: win-rate plots showing the percentage of datasets where each method achieves rank #1. GRNFormer is the top performer in 63% of datasets for AUROC and 50% for AUPRC. These plots reinforce GRNFormer's superior and consistent performance across evaluation settings.

#### C. Early Precision Calculation

To further evaluate early-ranking performance, we assessed Early Precision (EP), which quantifies how accurately a method prioritizes true regulatory interactions among the highest-confidence predicted edges. This metric focuses on the top-ranked predictions that are most biologically actionable and complements global measures such as AUROC and AUPRC.

For a given method, Early Precision at rank K is defined as:

$$\mathrm{EP@K} = \frac{\text{Number of true regulatory edges among top } K \text{ predictions}}{K}$$

Thus, EP@K directly measures the proportion of correct regulatory interactions within the top K predicted edges. Higher EP values indicate stronger prioritization of biologically valid regulatory interactions in the early ranking regime.

For each method and dataset configuration, all gene–gene edges were ranked according to their predicted confidence scores, after which the top K edges were selected for K $\in$ {100, 500, 1000, 2000, |GT|}. The overlap between these predicted edges and the corresponding gold-standard regulatory networks was then computed, and EP@K was calculated as the fraction of true positives among the top K predictions. This evaluation was performed consistently across non-cell-type-specific ChIP-seq networks, cell-type-specific ChIP-seq networks, and STRING functional interaction networks, as well as across all gene-set configurations (500, 1000, TF500, TF1000).

Supplementary Figure S10 presents Early Precision (EP) values for all evaluated methods across multiple ranking thresholds, including EP@100 (A), EP@500 (B), EP@1000 (C), EP@2000 (D), and EP evaluated at the number of ground-truth interactions (E), across non–cell-type-specific ChIP-seq networks, cell-type-specific ChIP-seq networks, and STRING interaction networks under all gene-set configurations.

Across non–cell-type-specific ChIP-seq and STRING benchmarks, EP values remain close to zero for most methods, indicating limited early recovery of true regulatory interactions in these broader regulatory settings. In contrast, meaningful early precision is observed primarily in the cell-type-specific ChIP-seq networks, where several learning-based approaches demonstrate the ability to prioritize true regulatory edges at early ranking depths.

At lower cutoffs (EP@100 and EP@gt), GRNFormer and GNE exhibit comparable early precision across many datasets, frequently alternating as the top-performing method depending on the regulatory context and gene-set configuration. GNNLink shows moderate performance in selected settings but with greater variability.

As the ranking depth increases to EP@1000 and EP@2000, GRNFormer consistently maintains higher EP values across most cell-type-specific benchmarks, than the half of other competing methods. This trend is further reinforced when EP is evaluated at the full set of gold-standard interactions (panel E), where GRNFormer emerges as one of the strongest ,with stable recovery of true regulatory edges across regulatory contexts.

To quantify relative performance across datasets, Supplementary Figure S11A reports the **win rate**, defined as the fraction of datasets in which each method achieved the highest EP at a given cutoff. GRNFormer frequently has the higher win rate at all the thresholds compared to most of the methods, while standing as competitive as GNE for lower K values and K equals to ground truth interactions (win rate ~50% for EPR@100,and EPR@gt). Supplementary Figure S11B presents the **rank frequency**, showing how often each method appears at each rank across datasets. GRNFormer consistently occupies one of the top 3 ranks on average 84% of the times across all the EP cutoffs, further confirming its stability and consistent early recovery of true regulatory interactions.

Overall, these complementary analyses (EP heatmaps, win rate, and rank frequency) demonstrate that GRNFormer not only performs strongly in top-ranked edge recovery but also maintains robust and consistent prioritization of biologically relevant regulatory interactions across diverse benchmarks.

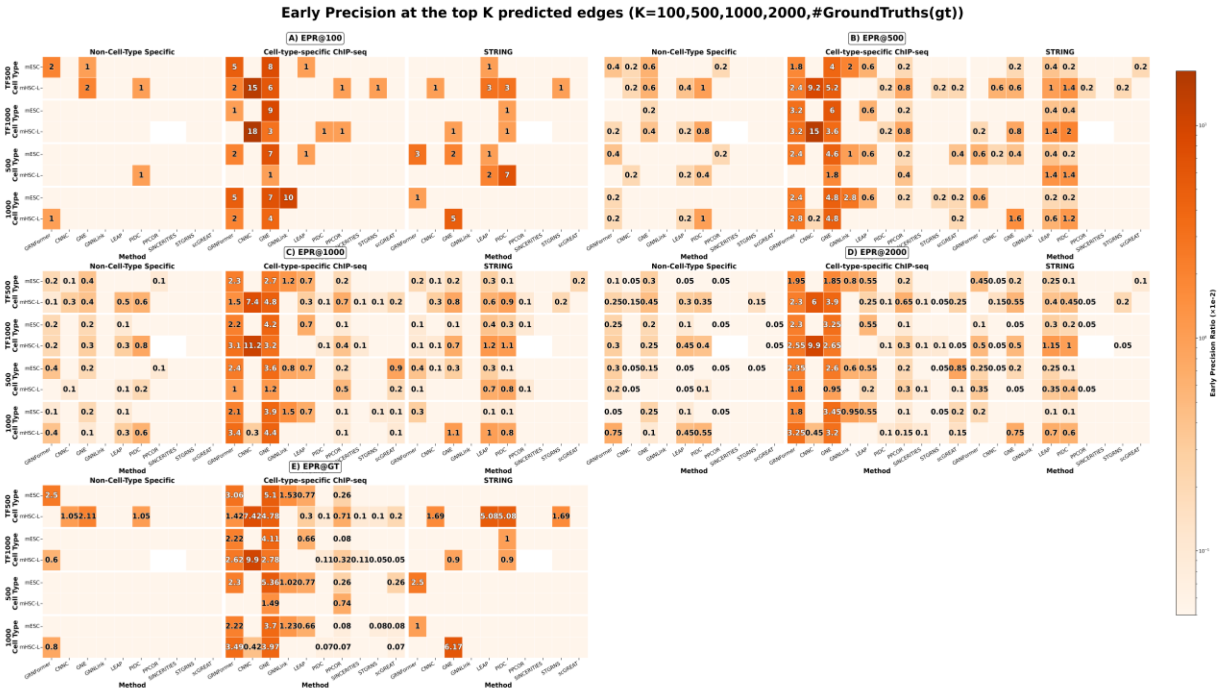


**Figure S10**. Early Precision (EP) values for all evaluated methods across multiple ranking thresholds, including (A) EP@100, (B) EP@500, (C) EP@1000, (D) EP@2000, and (E) EP evaluated at the number of ground-truth interactions, across non–cell-type-specific ChIP-seq networks, cell-type-specific ChIP-seq networks, and STRING interaction networks under all gene-set configurations.

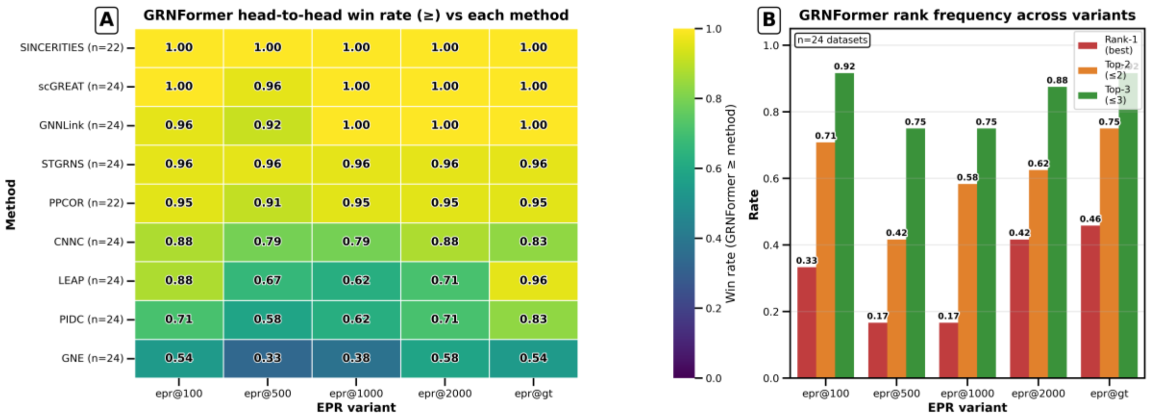


**Figure S11.A.** GRNFormer EPR Win rate vs each method (how often GRNFormer is top performing (⩾) than other methods at a given K). B. GRNFormer Rank frequency at each EPR Variant K.

#### S2. Detailed Ablation Study

Table S3 shows Ablation study details and corresponding performance. Removing the TF-Walker, which samples TF-anchored co-expression neighborhoods, and replacing with random sampling caused a substantial performance drop (Sampled_AUROC/ Sampled_AUPRC = 0.562/0.562), highlighting its critical role in focusing the model on biologically meaningful subgraphs. Replacing the Gene Transcoder with a linear embedding reduced contextual learning and led to a modest decline (Sampled_AUROC (SAUROC) = 0.931, Sampled_AUPRC (SAUPRC) = 0.906). Excluding the decoder, which contributes edge-level specificity, produced similar effects (Sampled_AUROC = 0.931, Sampled_AUPRC = 0.907), with further reduction when both components were removed.

Replacing the TransConv block with a Graph Convolutional Network (GCN) (Wu *et al.*, 2019) retained moderate performance (Sampled_AUROC = 0.928, Sampled_AUPRC =

0.904), but underperformed relative to the transformer-based architecture, indicating the value of pairwise attention for regulatory inference.

A minimal GCN-only baseline lacking all GRNFormer components yielded near-random results (Sampled_AUROC = 0.562, Sampled_AUPRC = 0.562), reinforcing that deep contextual modeling is essential for extracting regulatory signals from expression data.

Together, these results demonstrate that each GRNFormer module contributes distinct value: TF-Walker localizes transcriptional context, the Gene Transcoder and TransConv Block enrich representation learning, and the decoder sharpens edge inference. Their synergy enables accurate and generalizable GRN reconstruction across varied transcriptomic landscapes

#### S3. Cross-Species Generalization on DREAM5 Bulk RNA Seq Datasets

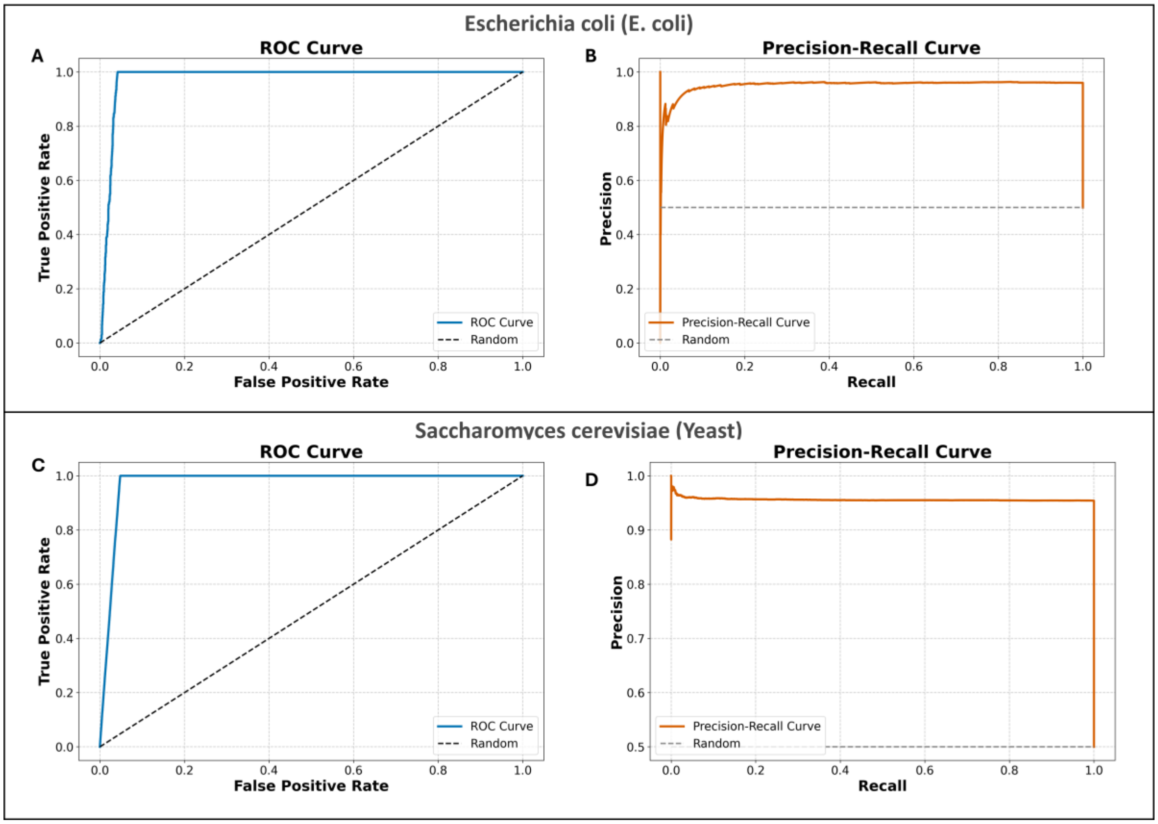


Figure S12: Blind evaluation of GRNFormer on DREAM5 bulk RNA-seq datasets for E. coli and S. cerevisiae and its scalability and cross-species generalization. (A) and (B). Sampled Precision-recall (PR) curve and sampled receiver operating characteristic (ROC) curve, respectively, for the E. coli gene regulatory network (GRN). (C) and (D). Sampled PR and sampled ROC curves for the S. cerevisiae GRN prediction.

#### S4. Perturbation Robustness

To assess the robustness of GRNFormer under perturbed single-cell expression conditions, we conducted a comprehensive evaluation by injecting two widely studied perturbations into the gene expression matrix prior to TF-Walker sampling:

1. **Scaled Gaussian noise** with standard deviation

   $\sigma \in \{0.1, 0.2, 0.3, 0.5\}$

2. **Random gene-expression dropout** applied independently with probability $p \in 0.1, 0.2, 0.3, 0.5$

For each $(\sigma, p)$pair, GRNFormer was run end-to-end using the same TF-Walker subgraphs and GCEN, and we measured AUROC and AUPRC relative to the unperturbed baseline

##### A. Experimental Design

For mHSC-L_non-cell-type-specific_chipseq_500 dataset, perturbations were applied directly to the expression matrix:

**Scaled Gaussian noise**

Noise was added *proportionally* to the magnitude of each expression value:

$$x'_{g,c} = x_{g,c}\,(1 + \mathcal{N}(0, \sigma^2)),$$

where $\sigma \in \{0.1, 0.2, 0.3, 0.5\}$.
This formulation introduces heteroscedastic perturbations similar to those observed in scRNA-seq technical noise.

**Random dropout**

With probability $p \in \{0.1, 0.2, 0.3, 0.5\}$, an expression value is replaced with zero:

$$x''_{g,c} = \begin{cases} 0, & \text{with probability } p, \\ x'_{g,c}, & \text{otherwise.} \end{cases}$$

**Normalization and downstream pipeline**

After perturbation, all subsequent steps follow TF-Walker inference module subgraph extraction, Gene transcoder embedding, GraViTAE encoding, and GRN inference, identically to the baseline model. Model performance was evaluated across **20 perturbation conditions** (4 noise levels × 4 dropout levels + unperturbed) using **three independent seeds**, providing stable and reproducible estimates of robustness.

#### B. Sampled_AUROC & Sampled_AUPRC Stability Under Perturbations

The perturbation curves are shown in Supplementary Fig. S13A & S13C. Across all tested perturbations, Sampled_AUROC remained remarkably stable. The maximum deviation from baseline was <0.003 (0.3%), indicating that GRNFormer's discriminative ability is essentially unaffected by substantial noise or sparsity. Even in the most challenging setting, where both dropout and scaled noise were simultaneously high ($\sigma$=0.5, p=0.5), Sampled_AUROC decreased only marginally from 0.9820 (baseline) to 0.9801. In several mild perturbation settings, Sampled_AUROC slightly exceeded baseline due to noise-driven amplification of weak but informative correlations—an effect well documented in high-dimensional regression. The perturbation curves exhibit smooth trajectories with no irregular fluctuations, further underscoring the stability of the model.

Sampled_AUPRC exhibited similarly robust behavior(Supplementary Fig. S13B & S13D). Across all perturbation combinations, the deviation remained below 0.0018 (0.18%), with baseline performance at 0.9668 and the worst-case setting reaching 0.9651. Since Sampled_AUPRC is particularly sensitive to false positives in sparse GRNs, this minimal change confirms that the internal TF-centered modeling performed by GRNFormer preserves regulatory precision even under pronounced noise and dropout. The observed variation falls well within the random-seed variability of the unperturbed system.

#### C. Architectural Basis for Robustness

The stability observed across all perturbation settings is not incidental; it emerges from three synergistic architectural characteristics of GRNFormer.

First, **z-score normalization** is applied across genes for each cell, which removes global scaling and mean shifts while preserving each gene's relative variation pattern. Because Pearson correlation is invariant to affine transformations, the normalization ensures that the GCEN-and therefore TF-Walker neighborhoods- remain nearly unchanged despite substantial perturbations.

Second, **TF-Walker** constructs TF-centered subgraphs based primarily on **correlation topology** rather than raw expression levels. As long as perturbations do not systematically distort correlation structure, the extracted neighborhoods remain highly consistent, allowing GraViTAE to receive comparable contextual information.

Third, **GraViTAE's variational latent representation** further mitigates sensitivity to noise. The encoder maps expression embeddings to latent Gaussian distributions, and the reparameterization sampling induces natural smoothing of perturbation-induced fluctuations. KL regularization suppresses overfitting to noise, and multi-head attention aggregates signals across multiple neighbors, diluting the influence of corrupted or dropped-out values. Together, these mechanisms enable the model to emphasize coherent co-expression patterns while suppressing spurious variation.

Supplementary Tables S5.1–S5.3 summarize the complete results. Aggregated Sampled_AUROC and Sampled_AUPRC values across all 20 perturbation settings show maximum degradation of <0.3% and <0.18%, respectively. Grouped summaries by noise level and dropout level confirm that performance remains stable even at the highest perturbation intensities.

Overall, the perturbation analysis on mHSC-L_non-cell-type-specific_chipseq_500 dataset demonstrates that GRNFormer maintains highly stable GRN inference accuracy under a wide range of noisy or sparse input conditions. This robustness reflects not only the capacity of the model but also its principled design: correlation-preserving preprocessing, context-consistent subgraph extraction, and variational smoothing through GraViTAE. These properties collectively explain why GRNFormer performs reliably across scRNA-seq datasets of varying quality, sparsity, and experimental noise profiles, further supporting its utility for cross-dataset and cross-species applications.

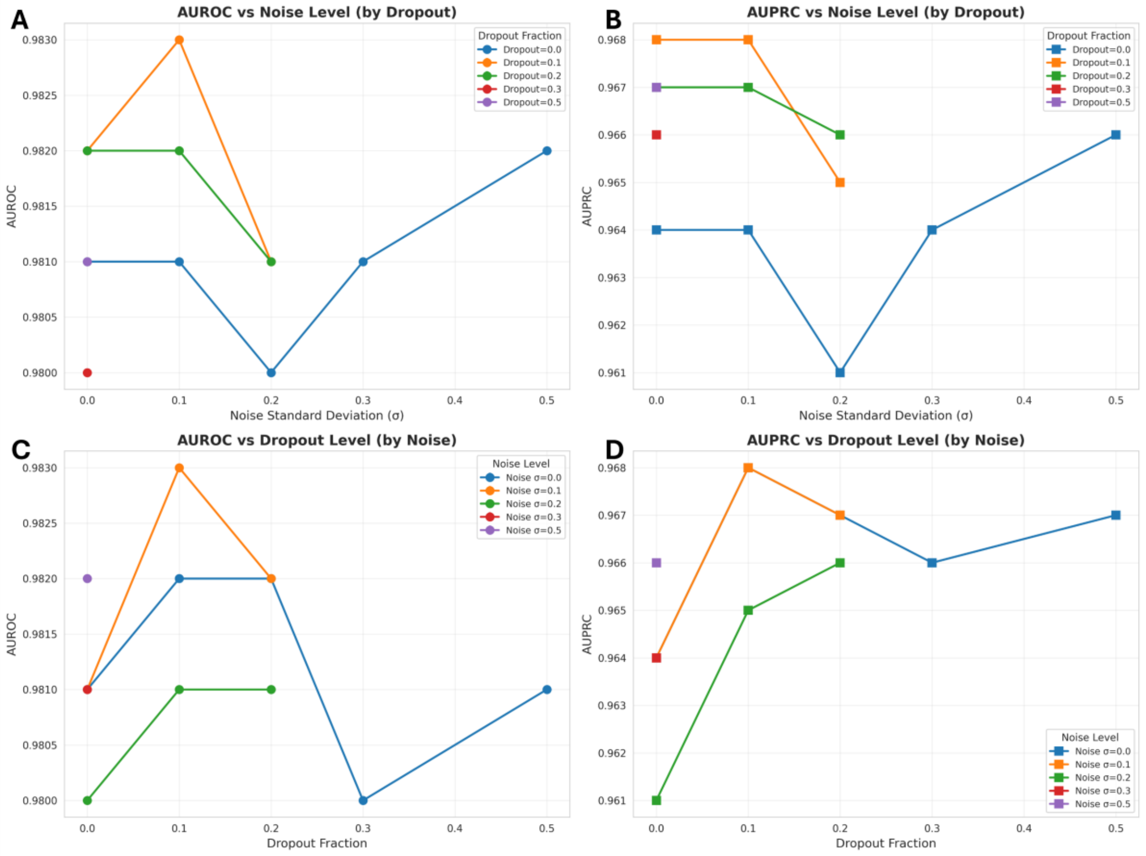


Figure S13. Robustness of GRNFormer to expression perturbations across combined Gaussian noise and dropout levels. (A) AUROC as a function of Gaussian noise (σ = 0.0–0.5), shown separately for each dropout fraction. (B) AUPRC under the same noise settings and dropout stratification.(C) AUROC as a function of dropout fraction (p = 0.0–0.5), stratified by noise level. (D) Corresponding AUPRC curves. Across all 20 perturbation combinations, performance degradation is minimal (ΔAUROC < 0.003; ΔAUPRC < 0.0018), indicating that GRNFormer remains stable even under substantial sparsity and noise. All AUROC and AUPRC are based on sampled evaluation.

#### S5. Biological Interpretation of Inferred Regulatory Links in hESC

The inferred network shown in Figure S14A, includes numerous interactions that align with known stem cell biology. *SOX2* was predicted to regulate *GATA3, EGR2,* and *POU2F1*, consistent with its role in modulating transcriptional plasticity and epithelial-mesenchymal transition during reprogramming (Nishimura *et al.*, 2017; Sarkar and Hochedlinger, 2013; Donner *et al.*, 2007). *MYC* was linked to cell cycle and biosynthetic regulators such as *TP53, NFYB*, and *FOXM1*, in line with its function in promoting proliferation and metabolic activity (Ahmadi *et al.*, 2021; Stine *et al.*, 2015). *POU5F1* was associated with *ETS2, ZBTB20,* and *PRDM1*, reinforcing its involvement in epiblast specification and developmental gene silencing (Gupta *et al.*, 2012; Fang *et al.*, 2018; Ermakova *et al.*, 2023). *NANOG*, another core regulator of naïve pluripotency, was connected to key early developmental factors including *NODAL, GDF3*, and *KLF2*(Papanayotou *et al.*, 2014; Duggal *et al.*, 2015; Mulas *et al.*, 2017). GRNFormer also captured several well-supported regulatory links validated by ChIP-seq data, including *SOX2-ZEB1, MYC-TCF7L1, MYC-SNAI1* and *NANOG-DKK1*, further confirming its ability to infer high-confidence regulatory edges from expression data alone (Herreros-Villanueva *et al.*, 2013; Morrison *et al.*, 2016; Baulida *et al.*, 2019; Zhu *et al.*, 2009). The targets of core pluripotency TFs (Figure S14A) were enriched in well-known stem cell pathways, including *TGF-β/SMAD* signaling (Attisano and Wrana, 2013), *Wnt* and *Hippo* signaling (Li *et al.*, 2019), circadian rhythm(Gao *et al.*, 2022), metabolic regulation(Burgess *et al.*, 2014), and chromatin remodeling(Martinez-Sarmiento *et al.*, 2024) (Figure S14B).

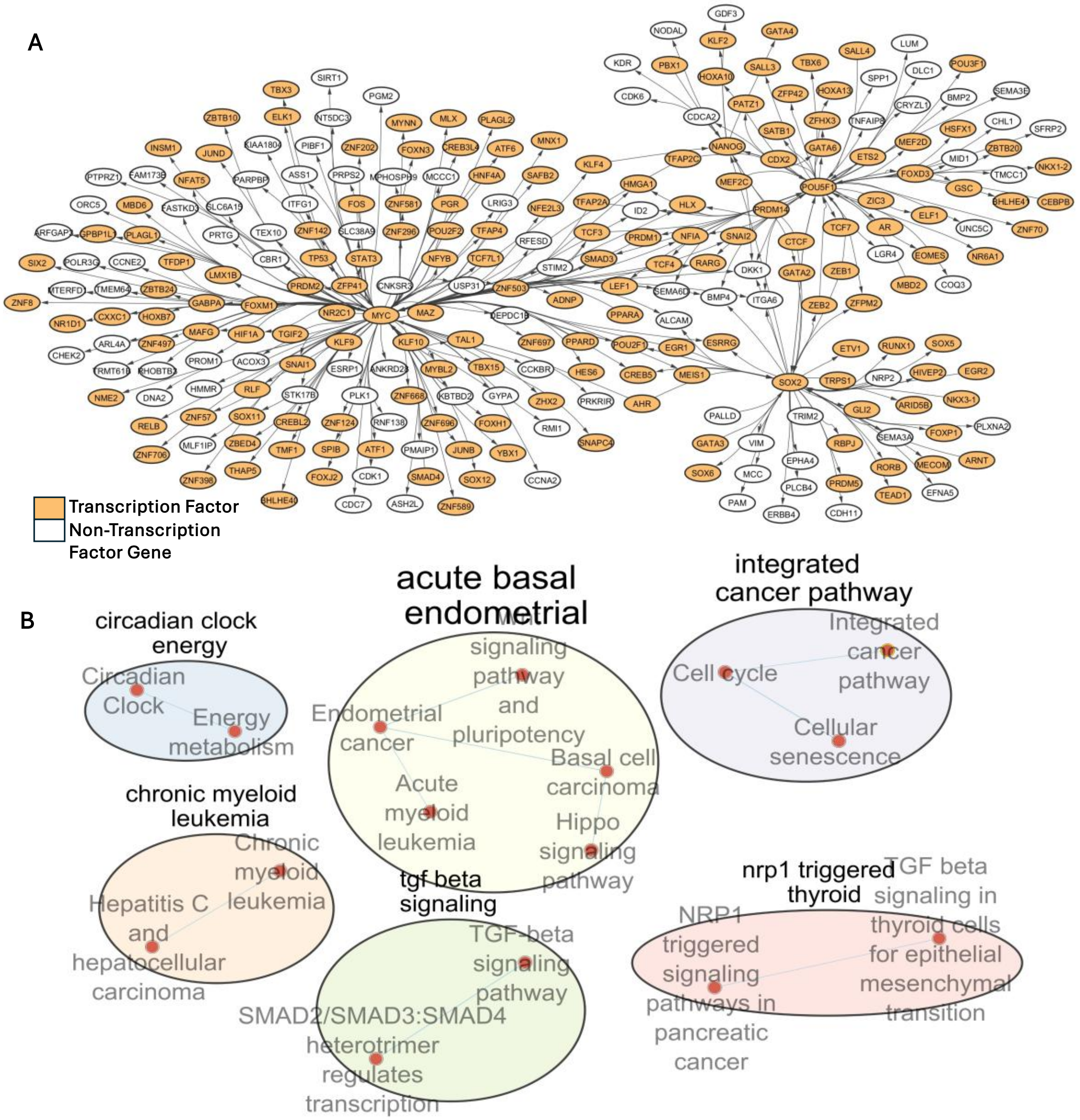


Figure S14: Application of GRNFormer to hESC cells. (A) GRNFormer predicted hESC-cell-type-specific GRN subgraph, centering around biomarkers of hESC cell types. The predicted GRN subgraph is verified against the ground truth network. (B) Enrichment map of GRN shown in S6A.

#### S6. Representation of Predicted GRN

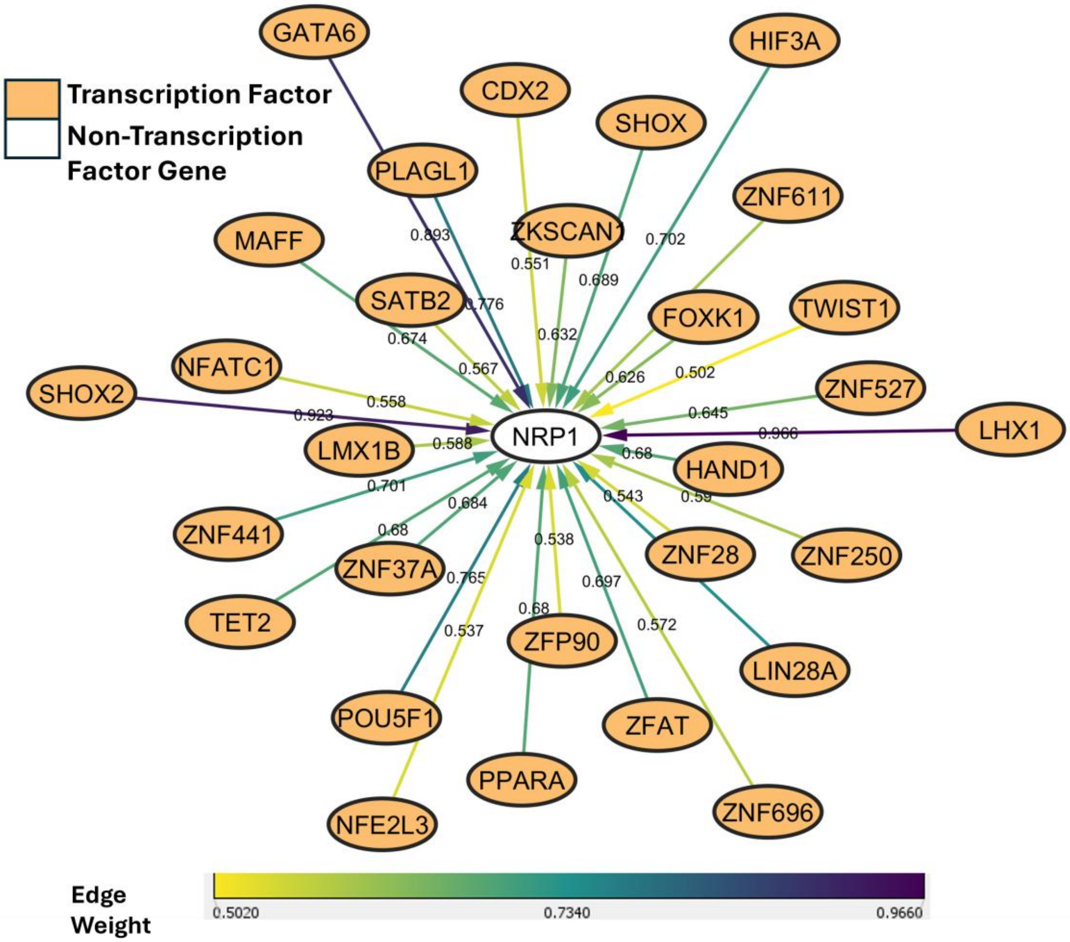


Figure S15: Predicted **NRP1**-centered subgraph (hESC). Nodes are genes; Edge color intensity ∝ GRNFormer predicted edge weight (low → high). Solid edges are novel predictions. The visualization highlights strong predicted associations between NRP1 and TFs such as POU5F1, LHX1, HAND1, GATA6, and TET2.

#### S7. Enrichment analysis Interpretation of Zero Short Regulatory Network Inference for PBMC

A central cluster shown in Figure S16B is generated from pathway enrichment analysis of network shown in Figure S16A, using g:Profiler and visualized the results in Cytoscape via Enrichment Map plugin. This cluster reflected general immune functions including vesicle trafficking, antigen presentation, and apoptotic signaling, suggesting basal regulatory programs across *T* cells and monocytes (Onnis *et al.*, 2016; Xu and Shi, 2007). A distinct cluster enriched for "graft-versus-host disease" and "viral process" mapped to monocytes and dendritic cells, known for their roles in antigen processing and cytokine signaling (Liu *et al.*, 2025; Marongiu *et al.*, 2021). Another module featured pathways like "DNA metabolic process," "complement activation," and "nucleotide excision repair," consistent with transcriptional activity in activated B cells and plasma cells (Dunkelberger and Song, 2010; Spivak, 2015).

Additional pathways such as "T cell migration," "immune effector regulation," and "leukocyte-mediated cytotoxicity" mapped to $CD8^{+}$ *T* cells and *NK* cells, reflecting cytotoxic immune functions (Rosenberg and Huang, 2018; Prager and Watzl, 2019; Uzhachenko and Shanker, 2019). Interaction module around *TAL1* inferred by GRNFormer is supported by literature evidence, that *TAL1* is a master regulator of hematopoiesis (Sanda *et al.*, 2012).

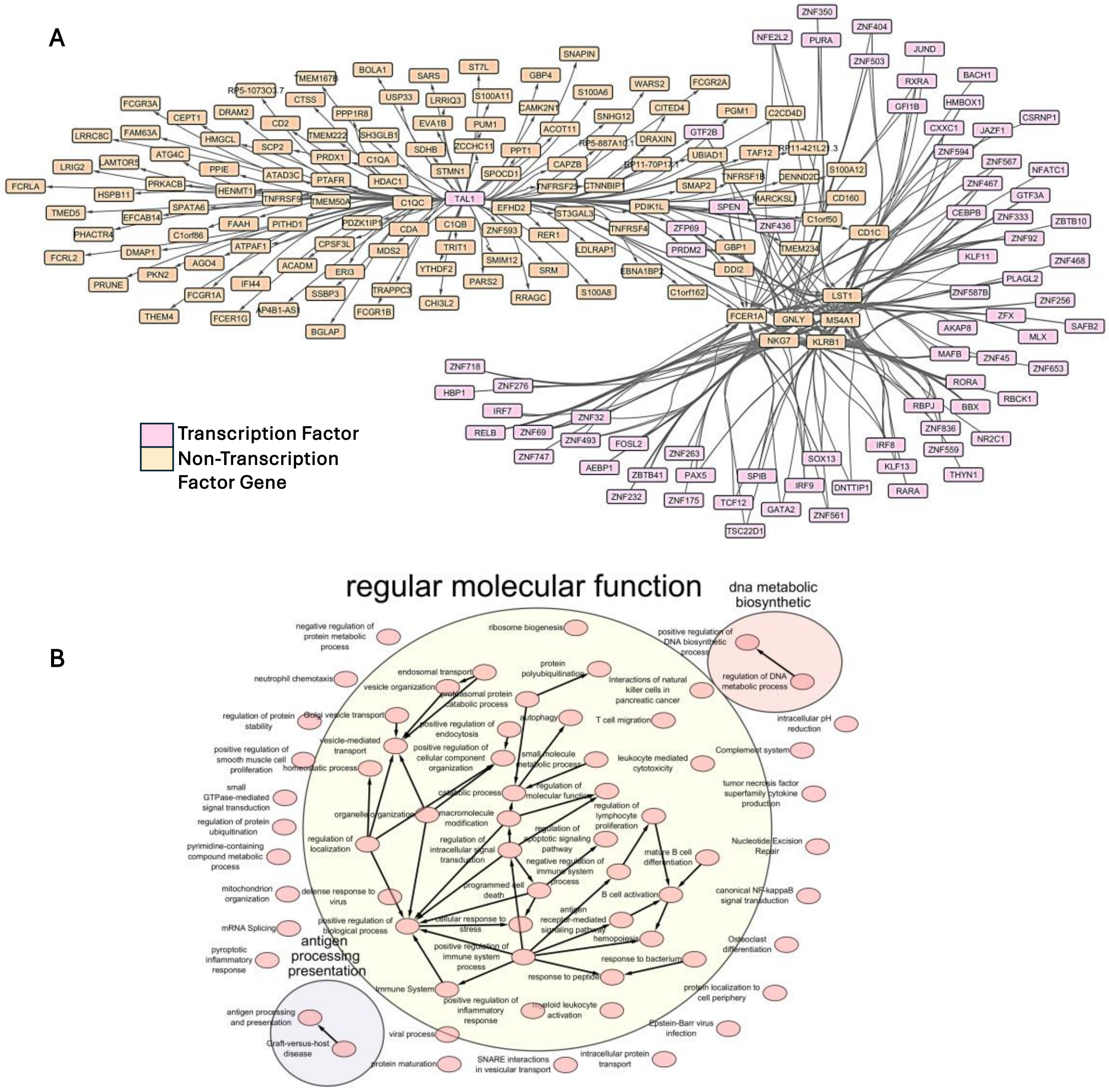


Figure S16: GRNFormer recovers cell-type-specific regulatory networks and functional modules from PBMCs via blind inference. (A) Subnetwork from the GRNFormer-predicted GRN on the PBMC 3k dataset, highlighting six key regulators (MS4A1, GNLY, KLRB1, LST1, FCER1A, and TAL1) corresponding to major immune lineages. The pretrained GRNFormer was applied to the dataset blindly without access to cell-type labels or prior pathway information. (B) Pathway enrichment map derived from GRN targets in (A), using g: Profiler and visualized with Enrichment Map in Cytoscape. Clusters represent immune processes such as antigen presentation, interferon signaling, cytotoxic activity, and B-cell DNA repair. Several predicted edges are supported by prior literature, validating the model's biological relevance.

#### Supplementary Tables

**Table S1: Detailed breakdown of performance of GRNFormer across five cell types across different regulatory contexts in the internal evaluation.** SAUROC and SAUPRC stand for Sampled_AUROC and Sampled_AUPRC. All the metric calculations shown here are based on bootstrapped sampled evaluation .

| Network Type | Cell Type | Gene Sets | #Genes | #True Positives | SAUROC | SAUPRC | Precision | Recall | F1 | Accuracy |
|---|---|---|---|---|---|---|---|---|---|---|
| Non-cell-type-specific ChIP-Seq | hESC | 1000 | 1000 | 649 | 0.955 | 0.913 | 0.92 | 0.841 | 0.879 | 0.884 |
| | | 500 | 500 | 181 | 0.96 | 0.929 | 0.933 | 0.89 | 0.911 | 0.913 |
| | | TF+1000 | 1697 | 8858 | 0.797 | 0.696 | 0.706 | 0.407 | 0.516 | 0.619 |
| | | TF+500 | 1197 | 6657 | 0.705 | 0.611 | 0.621 | 0.429 | 0.507 | 0.584 |
| | hHep | 1000 | 1000 | 207 | 0.98 | 0.96 | 0.967 | 1 | 0.983 | 0.983 |
| | | 500 | 500 | 68 | 0.977 | 0.958 | 0.962 | 1 | 0.981 | 0.98 |
| | | TF+1000 | 1559 | 7457 | 0.827 | 0.746 | 0.744 | 0.75 | 0.747 | 0.746 |
| | | TF+500 | 1059 | 5546 | 0.744 | 0.665 | 0.662 | 0.764 | 0.709 | 0.687 |
| | mDC | 1000 | 1000 | 460 | 0.976 | 0.956 | 0.954 | 1 | 0.977 | 0.976 |
| | | 500 | 500 | 34 | 0.984 | 0.973 | 0.966 | 1 | 0.982 | 0.982 |
| | | TF+1000 | 1588 | 8043 | 0.814 | 0.721 | 0.729 | 0.904 | 0.807 | 0.784 |
| | | TF+500 | 1088 | 6816 | 0.729 | 0.638 | 0.651 | 0.918 | 0.762 | 0.713 |
| | mHSC-E | 1000 | 1000 | 636 | 0.978 | 0.964 | 0.941 | 1 | 0.97 | 0.969 |
| | | 500 | 500 | 271 | 0.97 | 0.947 | 0.934 | 1 | 0.966 | 0.965 |
| | | TF+1000 | 1272 | 2174 | 0.934 | 0.906 | 0.848 | 1 | 0.918 | 0.91 |
| | | TF+500 | 772 | 1750 | 0.873 | 0.823 | 0.761 | 1 | 0.864 | 0.843 |
| | mHSC-GM | 1000 | 1000 | 454 | 0.984 | 0.976 | 0.95 | 1 | 0.974 | 0.973 |
| | | 500 | 500 | 122 | 0.981 | 0.966 | 0.956 | 1 | 0.978 | 0.977 |
| | | TF+1000 | 1206 | 1395 | 0.959 | 0.94 | 0.891 | 1 | 0.943 | 0.939 |
| | | TF+500 | 706 | 983 | 0.907 | 0.861 | 0.816 | 0.915 | 0.863 | 0.854 |
| Cell Type Specific ChIP-Seq | hESC | 1000 | 1000 | 2335 | 0.961 | 0.929 | 0.929 | 0.847 | 0.886 | 0.891 |
| | | 500 | 500 | 690 | 0.971 | 0.949 | 0.939 | 0.984 | 0.961 | 0.96 |
| | | TF+1000 | 1697 | 16742 | 0.816 | 0.73 | 0.735 | 0.439 | 0.55 | 0.641 |
| | | TF+500 | 1197 | 11590 | 0.736 | 0.656 | 0.664 | 0.512 | 0.578 | 0.626 |
| | hHep | 1000 | 1000 | 845 | 0.983 | 0.963 | 0.967 | 1 | 0.983 | 0.983 |
| | | TF+1000 | 1559 | 21240 | 0.848 | 0.77 | 0.762 | 0.805 | 0.783 | 0.777 |
| | | TF+500 | 1059 | 13809 | 0.774 | 0.694 | 0.684 | 0.802 | 0.739 | 0.716 |
| | mDC | 1000 | 1000 | 164 | 0.976 | 0.957 | 0.955 | 1 | 0.977 | 0.977 |
| | | TF+1000 | 1588 | 2128 | 0.807 | 0.725 | 0.714 | 0.846 | 0.775 | 0.754 |
| | | TF+500 | 1088 | 1444 | 0.722 | 0.642 | 0.639 | 0.873 | 0.738 | 0.689 |
| | mHSC-E | 1000 | 1000 | 4751 | 0.964 | 0.922 | 0.945 | 1 | 0.972 | 0.971 |
| | | 500 | 500 | 995 | 0.964 | 0.926 | 0.935 | 1 | 0.966 | 0.965 |

| | | | | | | | | | | |
|---|---|---|---|---|---|---|---|---|---|---|
| | | TF+1000 | 1272 | 22257 | 0.92 | 0.863 | 0.855 | 1 | 0.922 | 0.915 |
| | | TF+500 | 772 | 13854 | 0.862 | 0.788 | 0.768 | 1 | 0.869 | 0.849 |
| | mHSC-GM | 1000 | 1000 | 4900 | 0.974 | 0.947 | 0.955 | 1 | 0.977 | 0.976 |
| | | 500 | 500 | 1246 | 0.974 | 0.944 | 0.959 | 1 | 0.979 | 0.978 |
| | | TF+1000 | 1206 | 14821 | 0.945 | 0.899 | 0.898 | 0.795 | 0.843 | 0.852 |
| | | TF+500 | 706 | 9409 | 0.896 | 0.833 | 0.82 | 0.876 | 0.847 | 0.842 |
| STRING | hESC | 500 | 500 | 47 | 0.979 | 0.969 | 0.934 | 1 | 0.966 | 0.964 |
| | | 1000 | 1000 | 442 | 0.960 | 0.935 | 0.926 | 1 | 0.961 | 0.960 |
| | | TF+1000 | 1697 | 2732 | 0.815 | 0.742 | 0.736 | 0.49 | 0.588 | 0.657 |
| | | TF+500 | 1197 | 2446 | 0.731 | 0.663 | 0.653 | 0.457 | 0.537 | 0.607 |
| | hHep | 500 | 500 | 15 | 0.979 | 0.965 | 0.97 | 1 | 0.984 | 0.984 |
| | | 1000 | 1000 | 922 | 0.984 | 0.972 | 0.969 | 1 | 0.984 | 0.984 |
| | | TF+1000 | 1559 | 2666 | 0.847 | 0.798 | 0.752 | 0.782 | 0.767 | 0.762 |
| | | TF+500 | 1059 | 2200 | 0.772 | 0.729 | 0.67 | 0.799 | 0.729 | 0.703 |
| | mDC | 500 | 500 | 146 | 0.982 | 0.966 | 0.967 | 1 | 0.983 | 0.983 |
| | | 1000 | 1000 | 806 | 0.978 | 0.958 | 0.954 | 1 | 0.976 | 0.974 |
| | | TF+1000 | 1588 | 11422 | 0.829 | 0.758 | 0.732 | 0.914 | 0.813 | 0.79 |
| | | TF+500 | 1088 | 10037 | 0.751 | 0.677 | 0.654 | 0.928 | 0.767 | 0.718 |
| | mHSC-E | 500 | 500 | 276 | 0.977 | 0.963 | 0.934 | 1 | 0.966 | 0.965 |
| | | 1000 | 1000 | 701 | 0.982 | 0.972 | 0.942 | 1 | 0.970 | 0.969 |
| | | TF+1000 | 1272 | 2139 | 0.938 | 0.91 | 0.847 | 1 | 0.917 | 0.91 |
| | | TF+500 | 772 | 1765 | 0.881 | 0.838 | 0.762 | 1 | 0.865 | 0.844 |
| | mHSC-GM | 500 | 500 | 75 | 0.986 | 0.976 | 0.957 | 1 | 0.978 | 0.977 |
| | | 1000 | 1000 | 614 | 0.988 | 0.983 | 0.951 | 1 | 0.975 | 0.974 |
| | | TF+1000 | 1206 | 1451 | 0.961 | 0.946 | 0.89 | 1 | 0.942 | 0.938 |
| | | TF+500 | 706 | 960 | 0.907 | 0.867 | 0.818 | 0.912 | 0.863 | 0.855 |

#### Table S2: Overview of State-of-the-art GRN Inference Methods Compared with GRNFormer

| Method | Algorithmic Principle | Availability | Method Type |
|---|---|---|---|
| **LEAP** | Time-lagged correlation using lag-based Pearson correlation to infer directed edges. | R package (LEAP) | Statistical / Correlation-based |
| **PIDC** | Partial Information Decomposition of mutual information; accounts for redundancy/synergy. | Python (PIDC via ARACNE/IDTxl) | Information-theoretic |
| **PPCOR** | Partial correlation coefficient estimation via linear regression to infer conditional deps. | R package (ppcor) | Statistical / Correlation-based |
| **SINCERITIES** | Uses Granger causality and steady-state constraints via sparse linear ODE modeling. | https://github.com/Murali-group/Beeline.git | Dynamical System / ODE-based |
| **CNNC** | Convolutional neural network on 2D histogram of co-expression profiles. | https://github.com/xiaoyeye/CNNC.git | Deep Learning / Image-based |
| **GNNLink** | Graph Neural Network trained for edge prediction on constructed cell-gene graph. | https://github.com/sdesignates/GNNLink.git | GNN-based (Edge Prediction) |
| **GNE** | Learns node embeddings using skip-gram objective and predicts edges from embedding similarity. | https://github.com/kckishan/GNE.git | Embedding + Similarity (Node2Vec) |
| **scGREAT** | Transformer-based model using gene expression and gene bio text dictionaries | https://github.com/WangyuchenCS/scGREAT.git | GNN-based / Attention-based |
| **STGRNs** | Spatiotemporal transformer network modeling time-evolving GRNs from spatial data. | https://github.com/zhanglab-wbgcas/STGRNS.git | Spatiotemporal GNN / Attention |

**Table S3: Ablation study of GRNFormer components in blind cross-lineage evaluation. Each model was trained on five cell types (hESC, hHep, mDC, mHSC-E, and mHSC-GM), and blindly assessed on two new cell types (mESC and mHSC-L). AUROC and AUPRC averaged across the test sets of the two cell types are reported.** SAUROC and SAUPRC stand for Sampled_AUROC and Sampled_AUPRC. All AUROC and AUPRC calculations shown here are based on bootstrapped sampled evaluation .

| Ablation Variant | TransConv Block | Subgraph Sampling | Embedding Type | Decoder | SAUROC | SAUPRC |
|---|---|---|---|---|---|---|
| **Full GRNFormer (average)** | ✓ | **TF-Walker** | **Gene Transcoder** | ✓ | **0.970** | **0.950** |
| No TF-Walker | ✓ | Random Sampling | Gene Transcoder | ✓ | 0.562 | 0.562 |
| GCN (Full GCN-based variant) | ✗ (GCN) | TF-Walker | Gene Transcoder | ✗ | 0.928 | 0.904 |
| No Decoder (Edge Directly Predicted) | ✓ | TF-Walker | Gene Transcoder | ✗ | 0.931 | 0.907 |
| Simple Embedding Only | ✓ | TF-Walker | Linear Embedding | ✓ | 0.931 | 0.906 |
| No Gene Transcoder + No Decoder | ✓ | TF-Walker | Linear Embedding | ✗ | 0.930 | 0.906 |
| Plain Baseline (GCN only) | ✗ (GCN) | Random Sampling | Linear Embedding | ✗ | 0.562 | 0.562 |

**Table S4. GRNFormer performance statistics summary across subgraph sizes (10–500 nodes).** SAUROC and SAUPRC stand for Sampled_AUROC and Sampled_AUPRC. All AUROC and AUPRC calculations shown here are based on bootstrapped sampled evaluation.

| Dataset | Mean SAUROC ± SD | $CV_{AUROC}$(%) | Mean SAUPRC ± SD | $CV_{AUPRC}$ (%) | SAUROC Range | SAUPRC Range | # Sizes |
|---|---|---|---|---|---|---|---|
| **E. coli GRN** | **0.963 ± 0.022** | **2.28** | **0.942 ± 0.021** | **2.23** | **0.932–0.979** | **0.913–0.959** | **4** |
| **mESC ChIP-seq (TF500)** | **0.961 ± 0.007** | **0.77** | **0.930 ± 0.017** | **1.86** | **0.952–0.968** | **0.902–0.943** | **5** |
| **mESC ChIP-seq (TF1000)** | **0.963 ± 0.009** | **0.94** | **0.937 ± 0.011** | **1.18** | **0.947–0.969** | **0.918–0.946** | **5** |

**Table S5.1. Summary of GRNFormer Performance Across All Perturbation Conditions.** SAUROC and SAUPRC stand for Sampled_AUROC and Sampled_AUPRC. All AUROC and AUPRC calculations shown here are based on bootstrapped sampled evaluation.

| Perturbation Setting | SAUROC (mean) | SAUPRC (mean) | #count |
|---|---|---|---|
| **Baseline (no noise, no dropout)** | **0.9820** | **0.9668** | **3** |
| **Dropout-only (p ∈ {0.1, 0.2, 0.3, 0.5})** | **0.9818** | **0.9669** | **12** |
| **Noise-only (σ ∈ {0.1, 0.2, 0.3, 0.5})** | **0.9817** | **0.9652** | **12** |
| **Combined noise + dropout** | **0.9813** | **0.9654** | **12** |
| **Overall mean (all 20 perturbed conditions)** | **0.9816** | **0.9658** | **20** |

**Table S5.2. Performance Grouped by Dropout Level.** Performance varies by less than **0.3% AUROC** and **0.18% AUPRC** across dropout levels, showing high robustness to missing expression values. SAUROC and SAUPRC stand for Sampled_AUROC and Sampled_AUPRC. All AUROC and AUPRC calculations shown here are based on bootstrapped sampled evaluation.

| Dropout Fraction | SAUROC (mean) | SAUROC (std) | n | SAUPRC (mean) | SAUPRC (std) | #count |
|---|---|---|---|---|---|---|
| **0.0** | 0.9812 | 0.0008 | 5 | 0.9664 | 0.0015 | 5 |
| **0.1** | 0.9830 | 0.0017 | 3 | 0.9680 | 0.0006 | 3 |
| **0.2** | 0.9820 | 0.0010 | 3 | 0.9670 | 0.0006 | 3 |
| **0.3** | 0.9800 | – | 1 | 0.9660 | – | 1 |
| **0.5** | 0.9810 | – | 1 | 0.9670 | – | 1 |

**Table S5.3. Performance Grouped by Noise Level.** Even under strong perturbations (σ = 0.5), AUROC and AUPRC remain within **0.2–0.3%** of baseline, indicating that GRNFormer's predictions are stable even when substantial stochastic noise is injected. SAUROC and SAUPRC stand for Sampled_AUROC and Sampled_AUPRC. All AUROC and AUPRC calculations shown here are based on bootstrapped sampled evaluation.

| Noise σ | SAUROC (mean) | SAUROC (std) | n | SAUPRC (mean) | SAUPRC (std) | #count |
|---|---|---|---|---|---|---|
| **0.0** | 0.9812 | 0.0008 | 5 | 0.9664 | 0.0015 | 5 |
| **0.1** | 0.9820 | 0.0010 | 3 | 0.9663 | 0.0021 | 3 |
| **0.2** | 0.9807 | 0.0006 | 3 | 0.9640 | 0.0026 | 3 |
| **0.3** | 0.9810 | – | 1 | 0.9640 | – | 1 |
| **0.5** | 0.9820 | – | 1 | 0.9660 | – | 1 |

**Table S6. Statistical significance analysis of GRNFormer compared with GNE and GNNLink across all benchmark datasets.** Paired *t*-tests were conducted across 24 datasets to quantify whether GRNFormer's performance improvements over the strongest competing methods (GNE and GNNLink) are statistically significant under both evaluation regimes: bootstrap sampled evaluation and full-matrix evaluation. For each dataset, paired differences in AUROC, AUPRC, and AUPRC-ratio were computed, and the mean difference, win rate (percentage of datasets where GRNFormer outperforms the competing method), *p*-value, and significance level are reported. GRNFormer shows highly significant improvements over both competing methods in the sampled evaluation ($p < 0.001$ for all metrics), and maintains statistically significant gains under full-matrix evaluation, particularly against GNNLink. Significance thresholds: *** $p < 0.001$; ** $p < 0.01$; * $p < 0.05$; ns = not significant.

| Evaluation Type | Metric | Method | Mean Difference | Win Rate | p-value | Significance |
|---|---|---|---|---|---|---|
| Bootstrap Sampled | AUROC | GNE | 0.1485 | 91.7% (22/24) | $1.94\times10^{-5}$ | *** |
| Bootstrap Sampled | AUPRC | GNE | 0.130082 | 91.7% (22/24) | $7.79\times10^{-5}$ | *** |
| Bootstrap Sampled | AUROC | GNNLink | 0.1903 | 91.7% (22/24) | $3.40\times10^{-6}$ | *** |
| Bootstrap Sampled | AUPRC | GNNLink | 0.253299 | 91.7% (22/24) | $3.25\times10^{-10}$ | *** |
| Full-test-set | AUROC | GNE | 0.0708 | 70.8% (17/24) | 0.0283 | * |
| Full-test-set | AUPRC | GNE | 0.001170 | 70.8% (17/24) | 0.1120 | ns |
| Full-test-set | AUPRC-Ratio | GNE | 4.3509 | 70.8% (17/24) | 0.2221 | ns |
| Full-test-set | AUROC | GNNLink | 0.1078 | 83.3% (20/24) | $4.35\times10^{-4}$ | *** |
| Full-test-set | AUPRC | GNNLink | 0.004745 | 100.0% (24/24) | $8.94\times10^{-5}$ | *** |
| Full-test-set | AUPRC-Ratio | GNNLink | 12.1840 | 100.0% (24/24) | $2.78\times10^{-6}$ | *** |
